# *imply*: improving cell-type deconvolution accuracy using personalized reference profiles

**DOI:** 10.1101/2023.09.27.559579

**Authors:** Guanqun Meng, Yue Pan, Wen Tang, Lijun Zhang, Ying Cui, Fredrick R. Schumacher, Ming Wang, Rui Wang, Sijia He, Jeffrey Krischer, Qian Li, Hao Feng

## Abstract

Real-world clinical samples are often admixtures of signal mosaics from multiple pure cell types. Using computational tools, bulk transcriptomics can be deconvoluted to solve for the abundance of constituent cell types. However, existing deconvolution methods are conditioned on the assumption that the whole study population is served by a single reference panel, which ignores person-to-person heterogeneity. Here we present *imply*, a novel algorithm to deconvolute cell type proportions using personalized reference panels. *imply* can borrow information across repeatedly measured samples for each subject, and obtain precise cell type proportion estimations. Simulation studies demonstrate reduced bias in cell type abundance estimation compared with existing methods. Real data analyses on large longitudinal consortia show more realistic deconvolution results that align with biological facts. Our results suggest that disparities in cell type proportions are associated with several disease phenotypes in type 1 diabetes and Parkinson’s disease. Our proposed tool *imply* is available through the R/Bioconductor package *ISLET* at https://bioconductor.org/packages/ISLET/.

## INTRODUCTION

Tissues are complex samples composed of different cell types, and real bulk transcriptomic data are often weighted sums of multiple signals over several different cell types (Finotello and Trajanoski, 2018). In large-scale and population-level clinical studies, like Parkinson’s Disease Biomarkers Program (PDBP) and The Cancer Genome Atlas (TCGA), transcriptomic samples are often collected from complex tissues. For admixed tissue samples, differentially expressed transcriptional profiles from different phenotypical groups can be caused by either cell-type composition disparities or underlying cell-type-specific gene expression heterogeneity. Studies have shown that cell type proportions are confounders with other phenotypical covariates like age, sex, or clinical outcomes for bulk transcriptomic data analysis (Avila Cobos et al., 2020, 2018). As a result, ignoring cell-type-specific compositions in gene expression analysis would cause inflated false positive rates of identifying relevant genetic features. An accurate cell type proportion deconvolution is thus vital, especially for cell types with low abundance and weak biological signals where the real biological differences could be shadowed by technical noises (Kuhn et al., 2012; Avila Cobos et al., 2018; Meng et al., 2023).

Recently, several statistical methods have been proposed to deconvolute cell type abundance from bulk transcriptome data. These methods utilize the statistical framework of linear least squares regression (Tsoucas et al., 2019; Zhong et al., 2013; Clarke et al., 2010), quadratic programming (Gong and Szustakowski, 2013), support vector regression (Newman et al., 2015; Chiu et al., 2019), and non-negative matrix factorization (Gaujoux and Seoighe, 2012; Qiao et al., 2012). These methods share the same goal of quantifying the unknown abundances of various cell types and can be broadly summarized into two categories: Reference-Based (RB) and Reference-Free (RF). The RB deconvolution relies on a cell-type-specific gene expression signature reference panel composed of the pre-selected features known to differentiate cell types, while the RF deconvolution estimates cell type proportions in the absence of a reference panel. In general, reference-based approaches have better performance and lower error rates compared with reference-free approaches (Avila Cobos et al., 2020). Naturally, the accuracy of cell type abundance inference is dependent on the quality of signature matrices, and a more accurate reference panel is beneficial for improving cell type abundance estimations (Avila Cobos et al., 2020). RF deconvolution, in contrast, offers flexibility where reference panels are hard to obtain.

Currently, all RB deconvolution methods require a reference panel as the input across all subjects. For example, CIBERSORT(Newman et al., 2015), which is a state-of-art RB deconvolution approach (Avila Cobos et al., 2020, 2018), provides a verified signature panel LM22. It is specifically for leukocyte deconvolution and includes 547 marker genes which could distinguish 22 hematopoietic cell types. xCell (Aran et al., 2017) combines the gene set enrichment with deconvolution techniques and introduces curated gene signatures representing 64 distinct cell types, including a wide range of both adaptive and innate immune cells. However, it is a very strong assumption to use a single reference panel across the whole population. This assumption ignores person-to-person heterogeneity for cell-type-specific gene expression, and deviates from the biological fact that the gene expression profile could vary, even for one purified cell type, depending on environmental influences, age, sex, subject’s health status, and treatment paradigms (Kedlian et al., 2019; Di Biase et al., 2022; Findley et al., 2021; Idaghdour et al., 2010; Aguirre-Gamboa et al., 2016; Gibson, 2008; Çalişkan et al., 2015; Troester et al., 2004; Modlich et al., 2004). Mismatched reference signatures can impact the deconvolution results (Sutton et al., 2022; Ghaffari et al., 2023). The problem is even exacerbated when handling longitudinally observed and repeatedly-measured data, when intra-subject samples share information and inter-subject heterogeneities are relatively strong. Recent research shows that models incorporating personalized effects can accurately retrieve cell type reference panels on the individual-basis (Feng et al., 2023). However, to date, no method is available to take advantage of personalized references panel to precisely deconvolute cell type proportions, especially when longitudinal samples are available.

Here we develop a new deconvolution algorithm ***imply*** (***imp****roving ce****l****l-t****y****pe deconvolution using personalized reference*) as depicted in **Figure 1. *imply*** can utilize personalized reference panels to precisely deconvolute cell type proportions using longitudinal or repeatedly measured data. It borrows information across the repeatedly measured transcriptome samples within each subject, to recover personalized reference panels. The personalized references are further adopted to improve cell type deconvolution. The method consists of three stages. In the first stage, using a commonly shared reference panel across the population, we deconvolute the bulk transcriptomic data and estimate initial cell type proportions. The first stage method is based on support vector regression, as it has been shown to be a leading framework for conventional deconvolution problems (Newman et al., 2015). In the second stage, we use a mixed-effect modeling framework to retrieve personalized reference panels based on subjects’ phenotypical information, observed bulk transcriptomic data, and the initial cell type proportions from the first stage. In the third and final stage, we use the recovered personalized reference panels, together with repeated measurement of bulk transcriptomic data for each subject, to estimate cell type proportions. The rationale for using this three-stage approach is straightforward: the personalized reference panel is more accurate compared with the population-level signature. Naturally, using this more accurate reference panel can consequently lead to a more precise cell-type deconvolution.

**Figure 1.**
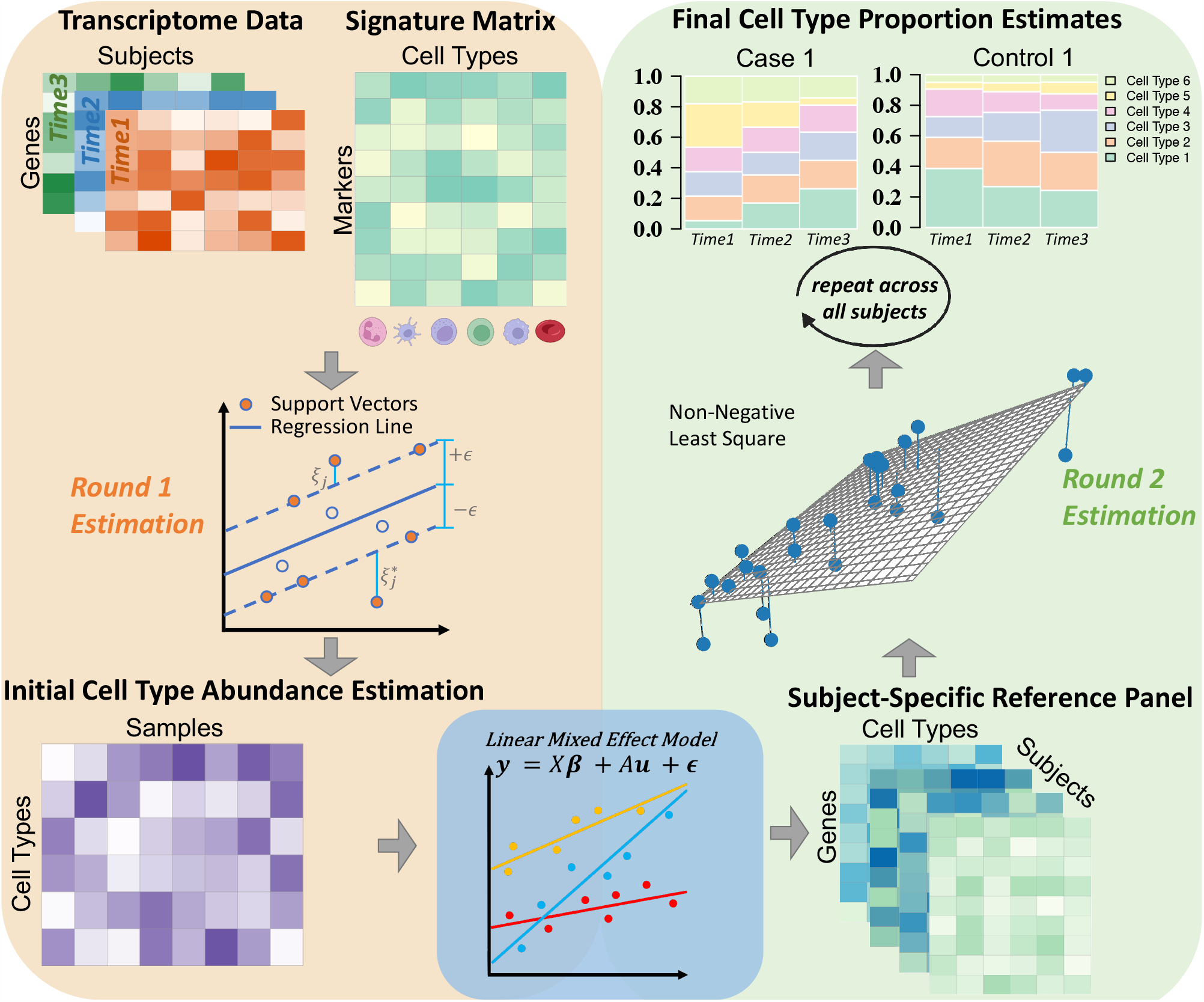
Overview of ***imply*** ‘s personalized deconvolution. The **top-left** shows two inputs: repetitively measured transcriptome data and a signature matrix containing cell-type-specific marker genes. In *Stage I*, depicted in the **middle left**, the initial step adopts support vector regression to derive a preliminary cell type abundance, as shown in the **bottom-left**. Next, for *Stage II*, as shown in the **bottom-center**, linear mixed effect models are utilized to reconstruct personalized references, which are shown in the **bottom-right**. In *Stage III*, as illustrated in the **middle-right**, by employing *non-negative least square* and using personalized references generated from the previous step, repeatedly across all subjects, ***imply*** enables personalized deconvolution to produce cell type proportion estimates, shown on the **top-right**.

We conducted extensive *in silico* simulations and real data analyses to test the performance of ***imply***. The simulation results showed significantly increased accuracies in cell type proportion estimation compared with existing approaches. Our method ***imply*** reduced bias in deconvolution, and increased the correlation between the estimated and the ground-truth cell type abundance. Real data analyses on two large longitudinal consortia, The Environmental Determinants of Diabetes in the Young (TEDDY study) and Parkinson’s Disease Biomarkers Program (PDBP study), showed more realistic deconvolution results that align with low-throughput experiments. The results suggested that disparities in cell type proportions of certain cell types are associated with several disease phenotypes in type 1 diabetes and Parkinson’s disease. Our method ***imply*** has been implemented and integrated into the Bioconductor package *ISLET* and is available at https://bioconductor.org/packages/ISLET/.

## MATERIALS AND METHODS

### Methods

#### Overview of imply

To outline briefly, the primary objective of ***imply*** is to improve the accuracy of cell abundance estimations through the integration of subject- and cell-type specific reference panels, termed personalized references. The algorithm is structured into three stages. In *Stage I*, the initial cell proportion estimates will be obtained. The core component of ***imply*** lies in *Stage II*, where a personalized reference panel is retrived for each subject. These personalized references will replace the population-level signature matrix, facilitating a personalized deconvolution process repeatedly across all subjects in *Stage III*.

#### Notation introduction

Let *G*(*g* = 1, 2, 3, *G*) denotes the total number of features (e.g., genes), and *N* (*n* = 1, 2, 3, *N* ) as the total number of subjects. For each subject *n*, there are *t*_*n*_(*i* = 1, 2, *t*_*n*_) repeated or longitudinal samples. The total number of samples across *N* subjects is thus 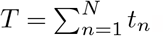 The bulk transcriptome dataset can be represented as a matrix ***Y***, of dimension G × T.

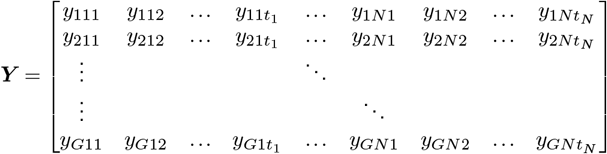

We denote *K*(*k* = 1, 2, … *K*) as the total number of purified cell types. Initially, we would have a population-level signature matrix ***E*** of dimension *J × K*(*J < G*), where *J* indicates the total number of discriminative signature genes for the first-round cell type deconvolution. This signature matrix, or reference panel, can be derived from pure cell line data or aggregated from annotated single-cell RNA-seq (scRNA-seq) data (Wang et al., 2019; Huang et al., 2023; Dong et al., 2021).

#### Stage I: Initial cell type proportion estimation

With the observed admixed data ***Y*** and the initial reference panel ***E***, as illustrated in the top-left of **Figure 1**, the first-round reference-based coarse deconvolution is conducted using a ν-Support Vector Regression algorithm (ν-SVR) (Schölkopf et al., 2000) based on a linearity assumption (Kuhn et al., 2011; Shen-Orr et al., 2010). Such strategy was already proven to be a successful choice in state-of-art deconvolution algorithms such as CIBERSORT (Newman et al., 2015). Support vectors are regulated by ν:-tubes integrated into the objective function (specified by the equation below).

To be specific, the algorithm in CIBERSORT requires both signature matrix ***E*** and reduced sample-specific RNA-sequencing data ***Y***, comprising only the overlapped features filtered based on marker genes from the signature matrix. The deconvolution is thus a regression problem: ***y*** _*ni*_ = *f* (***θ***_*E,ni*·_) = ***Eθ***_*E,ni*·_ +***b***, where ***b*** ∈ *R*^*J*^ capture random bias, and we can minimize the following objective function (Newman et al., 2015):

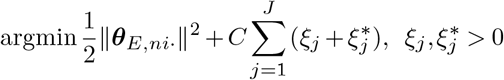

The solved 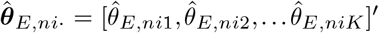 is the first-round sample-specific cell type abundance estimation. The constraints of the objective function and parameters of ϵ *C, ξ* _*j*_, and 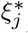 are detailed in the supplementary material section 1.1. Then negative coefficients (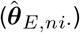) are set to 0, and the remaining coefficients are normalized to sum-to-one, which is the general practice in proportion deconvolution (Newman et al., 2015). Repeating this process for all samples, we obtain the deconvoluted cell composition matrix 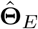 with dimension *T* × *K*. It’s worth to note that this first-step deconvolution of cell type proportions provides a valid initial estimation for downstream steps.

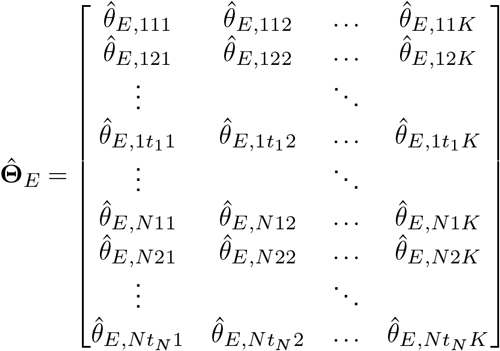

#### Stage II: Personalized reference panel recovery

The second step aims to retrieve a subject- and cell-type-specific reference panel. Using the cell-type-specific and sample-specific proportions 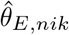 from stage I, we can set up the following linear mixed-effect regression for each gene *g*:

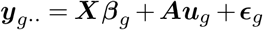

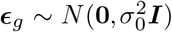 are the residuals and ***y***_*g*_ stands for the vector of observed expression data for a specific gene *g* for all *T* samples. Here ***X*** and ***A*** are the design matrices (each with dimension *T* × 2*K* and *T* × *NK*) for the fixed-effect ***β***_*g*_ and the random-effect ***u***_*g*_, respectively (details of ***X*** and ***A*** are specified in supplementary material section 1.2.) The initial cell type abundance information is further reorganized into vectors 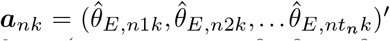 .The fixed-effect 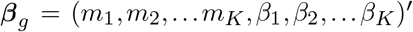 has two components: (*m*_1_, *m*_2_, … *m*_*K*_) are the baseline average cell-type-specific gene expression in the control group, and (*β*_1_, *β*_2_, … *β*_*K*_) are the ‘*difference*’ between the case group and the control group at cell type level. Note that our modeling allows for the incorporation of subject-level covariates such as disease status, for example, *z*_*n*_ = 1 for disease versus *z*_*n*_ = 0 for normal. The random-effect ***u***_*g*_ = (*u*_11_, *u*_21_, … *u*_*N*1_, *u*_12_, *u*_22_, … *u*_*N*2_, …, *u*_1*K*_, *u*_2*K*_, … *u*_*NK*_) ′ captures the subject-level and cell-type-specific gene expression deviation from the group-level average. 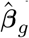 and û_*g*_ can be solved by penalized least square algorithm with restricted maximum likelihood (Bates et al., 2014). The subject- and cell-type-specific reference panel is obtained by combining 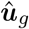 and û_*g*_ (fixed effect + random effect), with respect to each corresponding condition, cell type, and subject. We further reorganize reference panel into a subject-level structure and use ***R***_*n*_ with dimension *G* × *K* as shown below to represent the subject- and cell-type-specific reference panel. Each element in ***R***_*n*_ could be retrieved as: 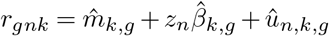

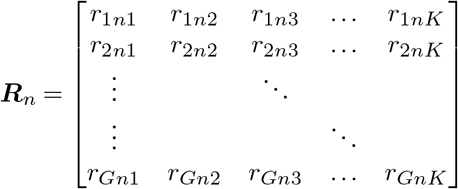

#### Stage III: Personalized deconvolution

With the subject-specific reference panel ***R***_*n*_ available for each subject *n*, and the original bulk mixture transcriptome data, as shown in the lower-right corner of **Figure 1**, we use *non-negative least square* (Lawson and Hanson, 1995, 1974) to deconvolute the cell type abundance **Θ**_*I,n*_. **Θ**_*I,n*_ is of dimension *K* × *t*_*n*_ for each subject respectively and the *I* in the subscript stands for the ***imply*** -estimated cell type abundance. To be specific, we optimize the following objective function in *non-negative least square*:

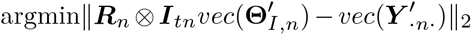

under the constraint **Θ**_*I,n*_ ≥ 0. ***Y*** _·*n*·_ is the bulk mixture data for subject *n* with dimension *G* × *t*_*n*_. This is a joint optimization across all the samples per subject simultaneously instead of sample-wise optimization, using the subject-specific signature matrix ***R***_*n*_ and quadratic programming. Note that ν -SVR with sample-wise optimization can also be utilized in this stage as an alternative approach, similar to *stage I*, and we name it ***imply-s***. Overall, instead of using the population-level signature matrix ***E***, the adoption of personalized ***R***_*n*_’s, for more genes, would consequently benefit cell type abundance inferences.

### Simulations

#### Pure cell-type-specific expression profiles

Notations of gene *g*, subject *n*, sample *i*, and cell type *k* are the same as the *Methods* section. The simulation scheme is borrowed and adapted from on our prior benchmark study (Meng et al., 2023), offering a comprehensive and flexible simulation framework. We utilized a set of true cell line RNA-seq dataset (Linsley et al., 2014) to obtain the distribution of gene expression parameters in a genome-wide scale. This study has six immune cell types (neutrophils, monocytes, B-cells, CD4 T cells, CD8 T cells, and natural killer cells). For each cell type, the cell-type-specific gene expression parameters, expression means (*μ*_*gk*_) and biological dispersion (*ϕ*_*gk*_), are obtained by using the *PROPER* (Wu et al., 2015) package. There are correlations across cell type for both expression means and dispersion, as expected. Therefore, for the reference panel simulation, we use *Multivariate Normal Distribution* (MVN) to capture correlations for both expression mean and dispersion, in the log scale. We use 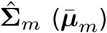 and 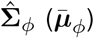 to de-note variance-covariance matrices of expression mean and dispersion, respectively. The dimensions match the number of cell types and the details of 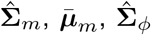 and 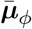 can be found in the supplementary materials section 2.1. We conduct 30 iterations for each simulation scenario, with six cell types and 1,000 genes:

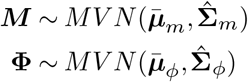

Note that the mean expression ***M***, and the biological dispersion **ϕ** are still parameter matrices for down-stream usage. The case and control groups share the same **ϕ**, but distinct mean expressions. The effect size of differential expression is defined by Log-Fold-Changes (LFC) denoted by Δ. The means for control and case are denoted by ***M*** _*Ctrl*_ = ***M*** and ***M*** _*Case*_ = ***M*** + Δ. We introduce 10% of differentially expressed (DE) genes on cell types 1, 2, 4, and 5, respectively. The true cell-type-specific gene expression matrix ***P*** is derived from a *Gamma Distribution* for both case and control:

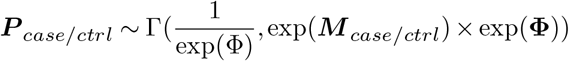

Subject-to-subject variations (SSV) are also introduced, implemented as the expression change percentages over the baseline in ***P*** _*case/ctrl*_. Variations are then added to ***P*** _*case/ctrl*_ to obtain subject-specific underlying gene expression matrices ***P*** _*n*_. To reflect the various levels of variations, the level of SSV can take the following ranges: 0-5%, 5%-10%, 10%-20%, and 20%- 50%. The total subject count per case/control group can take value in 25, 50, 75, and 100. The subject-level cell-type-specific underlying gene expression is shared across multiple samples, and each subject is measured 3 times.

#### Cell type proportions and observed read counts

To generate the cell type proportions, we borrow information from multiple well-labeled single cell RNA-seq studies. We mix and bootstrap cell labels from a combined pool and obtain the empirical cell proportions from this resampling. We use *Dirichlet Distribution* to estimate ***α*** parameters and simulate cell type proportions. The detailed procedures for generating cell proportions are outlined in supplementary materials section 2.2. The simulated sample-specific cell proportions are:

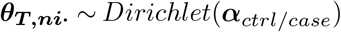

***θ***_***T***,***ni***_ are reorganized into cell composition matrix, **Θ**_*T*_ . The sample-specific underlying gene expression reference panel is the weighted average across cell types in ***P*** _*n*_ by ***θ***_*T,ni*·_, denoted as 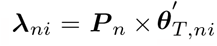, and will follow a *Gamma Distribution* as well (Moschopoulos, 1985). ***λ***_*ni*_ is further assessed by the *Poisson Distribution* to generate observed RNA-sequencing counts data, denoted as:

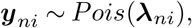

for subject *n* at measurement *i* across all *G* genes. Overall, the *Gamma Distribution* models biological variations, the *Dirichlet Distribution* regulates cell type proportion variations, and *Poisson Distribution* mimics technical noise related to the randomness in the sequencing experiments. This multi-step simulation design enables the separation of biological and technical noise (Meng et al., 2023; Feng et al., 2023), among other factors, to facilitate a comprehensive simulation study for our model testing.

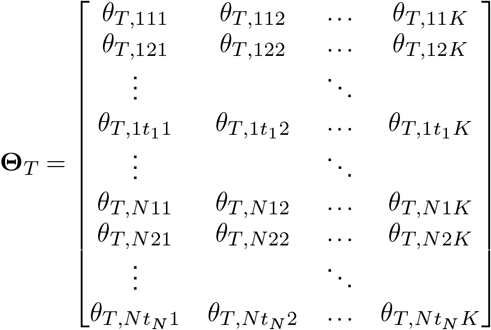

#### Input signature matrix

The signature matrix required by the algorithm as an input. To obtain it, we first take the average across all ***P*** _*n*_ matrices to get cell-type-specific gene expression mean matrix. Then 300 or 500 pseudo-marker genes are selected by *findRefinx* function (ordered by coefficients of variation) from TOAST (Li et al., 2019) to establish a signature matrix as the input for ***imply*** .

### Evaluation Metrics

We use **Θ** and 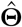 to denote the ground truth and estimated cellular abundances, which has the unique property of unit-sum and bounded by zero and one. Naturally, a central goal here is to assess how good the cellular abundances estimator 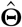 is. Specifically, we denote ***imply*** ‘s deconvolution values as 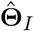, and existing method’s deconvolution results as 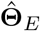 . The existing methods include currently available deconvolution approaches and those do not consider personalized reference panels. The following evaluation metrics are adopted for benchmarking:

#### 1. Absolute bias differences (ABD) and relative absolute bias differences (rABD)

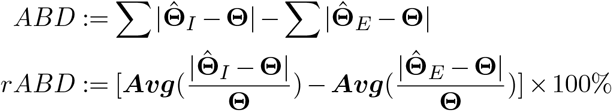

Here, for both *ABD* and *rABD*, if they are smaller than zero, it means the ***imply*** successfully reduces the estimation bias. A smaller value further indicates better performance.

#### 2. Correlation differences (CD)

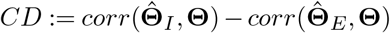

Here, if *CD>*0, then ***imply*** increases the correlation between the estimation and the ground truth. A larger value indicates favorable performance.

#### 3. Lin’s concordance correlation coefficient (CCC) and its variations

Lin’s concordance correlation coefficient (Lin’s CCC) (Lawrence and Lin, 1989), denoted as *ρ*_C_, has been extensively used to evaluate the concordance between a new measure and a gold standard measurement, and is defined as:

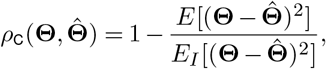

where *E*_*I*_ indicates the expectation under the as-sumption that **Θ** and 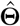 are independent. Lin’s CCC is bounded between 1 (perfect agreement) and -1 (disagreement), and the concordance improves as *ρ*_C_(**Θ**, 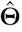 ) approaches 1. Additionally, we adopt a Euclidean distance-based variation of Lin’s CCC, by substituting the expected squared difference to Euclidean distance, denoted as *ρ*_C,E_, defined below:

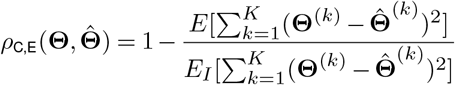

Another option is to employ the Aitchison (Aitchison et al., 2000) distance-based Concordance Correlation Coefficient (CCC), which is explained in detail in supplementary material section 1.3, with the results provided in supplementary material section 3.6. These metrics are adopted because they have been shown to be statistically more rigorous in dependent measures that are subject to the positiveness and unit-sum constraints (Cui et al., 2021), as is often the case in compositional proportion outcome. If ***imply*** yields increased concordance and improved precision, we would expect positive values in the differences of CCC. These metrics are respectively defined below:

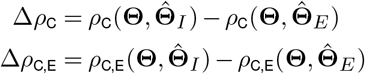

## RESULTS

We first evaluate ***imply*** ‘s deconvolution accuracy using synthetic data generated through the steps described earlier. ***imply*** is the only method that re-estimates cell type proportions using subject-specific reference panels from longitudinal bulk data; therefore, a direct comparison with existing deconvolution methods is not directly available. Nevertheless, we designed the benchmark to be inclusive of existing methods. TCA (Rahmani et al., 2019), designed for csDE genes detection, integrates a re-estimation feature for refining initially noisy cell proportion inputs. Specifically, TCA takes a maximum-likelihood (ML) approach to derive model parameters given initial cell proportion, and then the proportions are subsequently updated based on these estimated parameters. TCA requires preliminary cell proportions for effective re-estimation. We employ the *non-negative least squares* and *?*-SVR to acquire the initial inputs for TCA, and label them as TCA-n and TCA-s, respectively, which could be bench-marked with ***imply***. ISLET (Feng et al., 2023) is the first method to retrieve individual-specific reference estimation in repeated samples based on the Expectation-Maximization (EM) algorithm. ISLET can be an alternative approach to our mixed-effect model to solve subject-specific reference panels. Here, we consider ISLET-s and ISLET-n, respectively, representing ISLET variants that the final personalized deconvolution is conducted by SVR or *non-negative least squares*, respectively. We also introduce a variant of ***imply***, where *Stage III* is achieved by SVR instead of the *non-negative least squares*. This variant is denoted as ***imply-s***. We comprehensively benchmark our proposed personalized deconvolution methods, ***imply*** and its variant ***imply-s***, against other algorithms: TCA-n, TCA-s, ISLET-n, and ISLET-s.

### imply increases precision in cell-type deconvolution

We start with a baseline simulation scenario with six cell types, two disease groups, and 100 subjects per group with 3 replicates per subject. The subject-specific variation (SSV) in the underlying cell-type-specific gene expression panels is up to 5%. To simulate csDE genes, we introduce 10% of DE genes respectively to cell types 1, 2, 4, and 5. The effect size is characterized by the Log-Fold-Change (LFC) set to 0.5. **Figure 2A** shows the estimated reference panels by ***imply*** versus the ground truth. Overall, we observe good accuracy in personalized reference panel recovery, especially among high-expression genes. This result demonstrates the fidelity of *Stage II* and lays a foundation for personalized deconvolution in *Stage III*. Next, we evaluate if ***imply*** ‘s final cell type deconvolution, from *Stage III*, could reduce bias. Here, there are mainly two aspects to consider for accuracy benchmarking: one is to compare with alternative frameworks that do not use personal-ized reference panels; the other one is to benchmark with existing methods. **Figure 2B** shows the scatter-plot of the estimated cell type proportions versus the true proportions. Our result is overlaid on top of the result from CIBERSORT, one of the state-of-the-art methods. ***imply*** yields higher precision in deconvolution as its estimates aggregate closer to the diagonal line. In **Figure 2C-F**, the bias reductions are quantitatively assessed and compared using metrics introduced previously: *ABD, rABD, CD*, and Δ*ρ*_*C,E*_. Each point in a boxplot represents one simulation iteration, with the red dotted lines of zero indicating the basis for not using personalized reference panels. Thus, the zero line represents the existing deconvolution method, such as CIBERSORT, which did not consider personalized reference panels. For *ABD* and *rABD*, lower values indicate a greater increase in deconvolution accuracy; while for *CD* and Δ*ρ*_*C,E*_, higher values indicate improved concordance with the true values. Notably, ***imply*** consistently demonstrates the most substantial reduction in deconvolution bias and highest concordance with the truth. In contrast, TCA performs poorly, especially when the initial proportion inputs are estimated through *non-negative least squares* (TCA-n). Even when the initial proportion input is derived from CIBERSORT, the bias reduction achieved by TCA (TCA-s) is not as significant as that achieved by ***imply***. Furthermore, we notice that subject-specific reference panels estimated by ISLET also yield benefits for personalized deconvolution, illustrated by ISLET-s and ISLET-n. However, the improvements are not as pronounced as those achieved by ***imply***.

**Figure 2.**
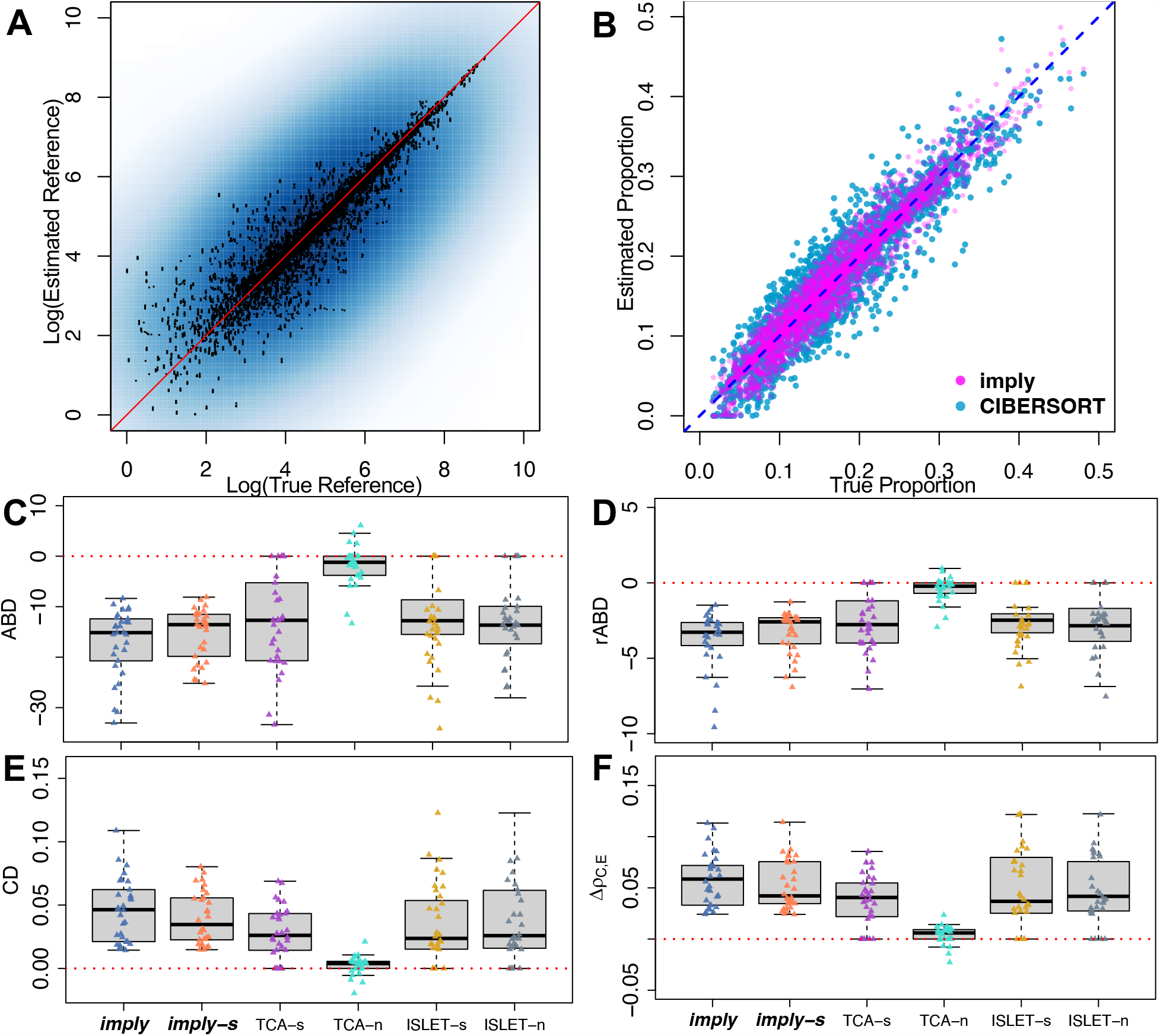
***imply*** can improve cell type deconvolution accuracy. **(A)** Scatterplot showing ***imply*** estimated gene expression reference panel versus the true reference panel values. **(B)** Superimposed scatterplot of the ***imply*** -estimated cell type proportion over the CIBERSORT-estimates, which are the results from the current state-of-art method. ***imply*** shows better concordance with the ground truth. **(C)**-**(F)** Boxplots displaying evaluation metrics and each point representing one simulation iteration: *ABD, rABD, CD*, and Δ*ρ*_C,E_. Five additional modeling frameworks are benchmarked. The red dashed line (value of 0) represents no improvement in proportion estimation. For **(C)** and **(D)**, lower values indicate better deconvolution accuracy. For **(E)** and **(F)**, higher the better.

We also explore the methods’ performance under various simulation scenarios and summarize the results in **Table 1**. The table shows averaged *ABD*s across simulation replicates, with each standard error, at exhaustive combinations of subject-specific variations (SSV=0-5%, 5%-10%, 10%-20%), effect sizes (LFC = 0.5, 1, 1.25), and sample sizes (N=25, 50, 100). Bold fonts highlight the algorithm with the most amount of bias reduction for each scenario. ***imply*** and ***imply-s*** consistently demonstrate exceptional performance in reducing deconvolution bias across all scenarios.

**Table 1.**
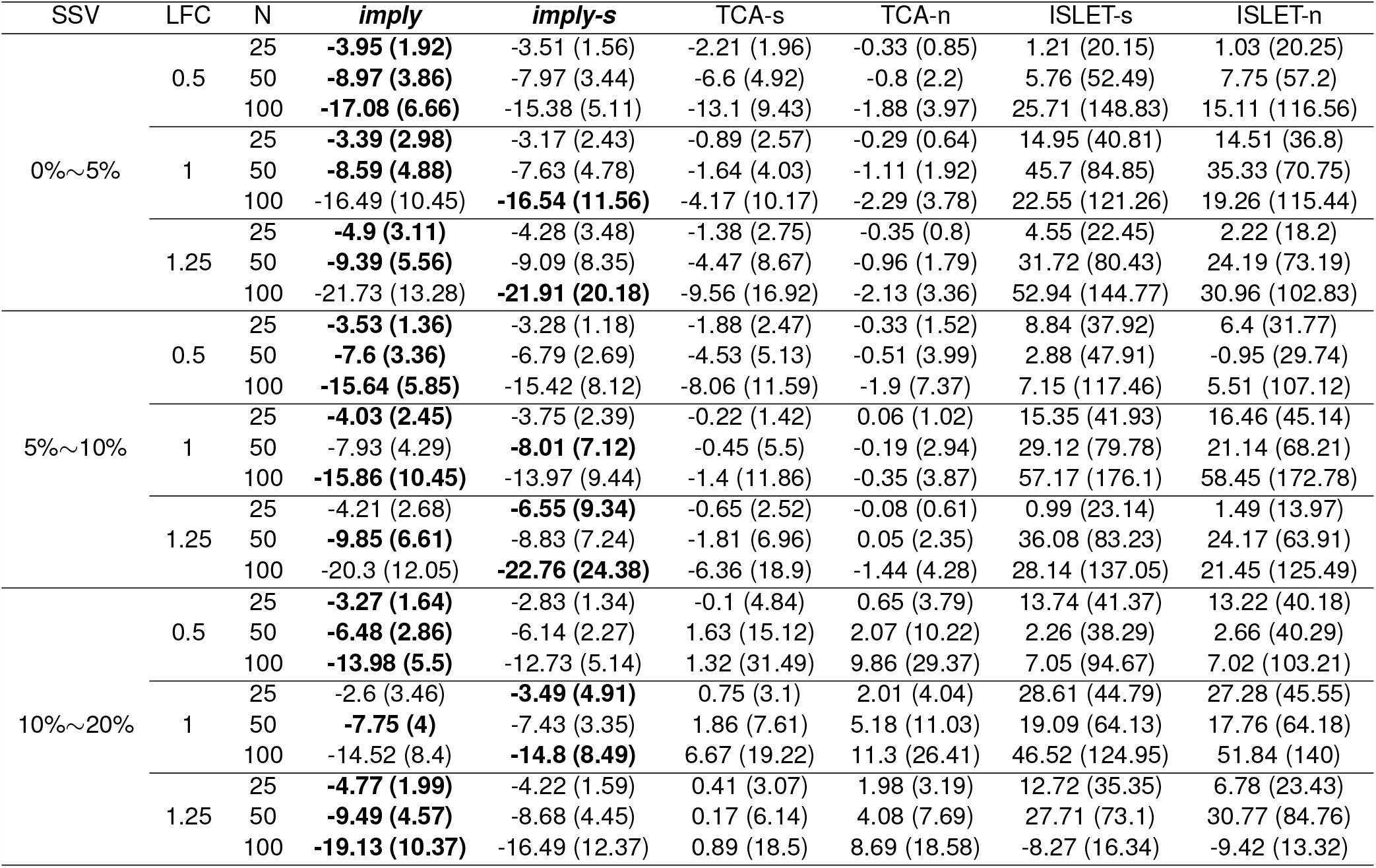
Benchmarking ***imply*** across various simulation scenarios. The table shows *Absolute Bias Difference (ABD)* at various subject-specific variations (SSV), effect sizes (LFC), and sample sizes (N). *ABD* values are shown, along with their standard error in parentheses. A lower value indicates better deconvolution estimation improvement. The bold font indicates the best method in each scenario.

### Benchmarking at cell-type resolution

We next investigate the deconvolution accuracy at each cell type. **Figure 3A** shows the *ABD* and Δ*ρ*_C_ outcomes of 30 replicates of each cell type under the condition of the SSV range of 0-5%, the sample size of 75, and the effect size of 0.5. Across all cell types, we can see a discernible reduction in bias when personalized reference panels are adopted. ***imply*** and ***imply-s*** consistently stand out, yielding a significant enhancement in concordance within each cell type compared to other models. The heatmap in **Figure 3B** shows the average *rABD* at various combinations of sample sizes and effect sizes, separated by cell types. At large effect sizes, improvements in cell deconvolution accuracies facilitated by ***imply*** are notably more profound. However, *rABD* exhibits limited alterations to variations in sample sizes. The simulation results also suggest a connection between bias reduction and cell type abundances; specifically, deconvolution accuracies for more abundant cells are highly sensitive to LFC changes (see supplementary materials section 3 for additional details). In contrast, for minor cell types, the small amount of contribution makes deconvolution an even more challenging task, where the sequencing noise could easily dominate underlying biological variations.

**Figure 3.**
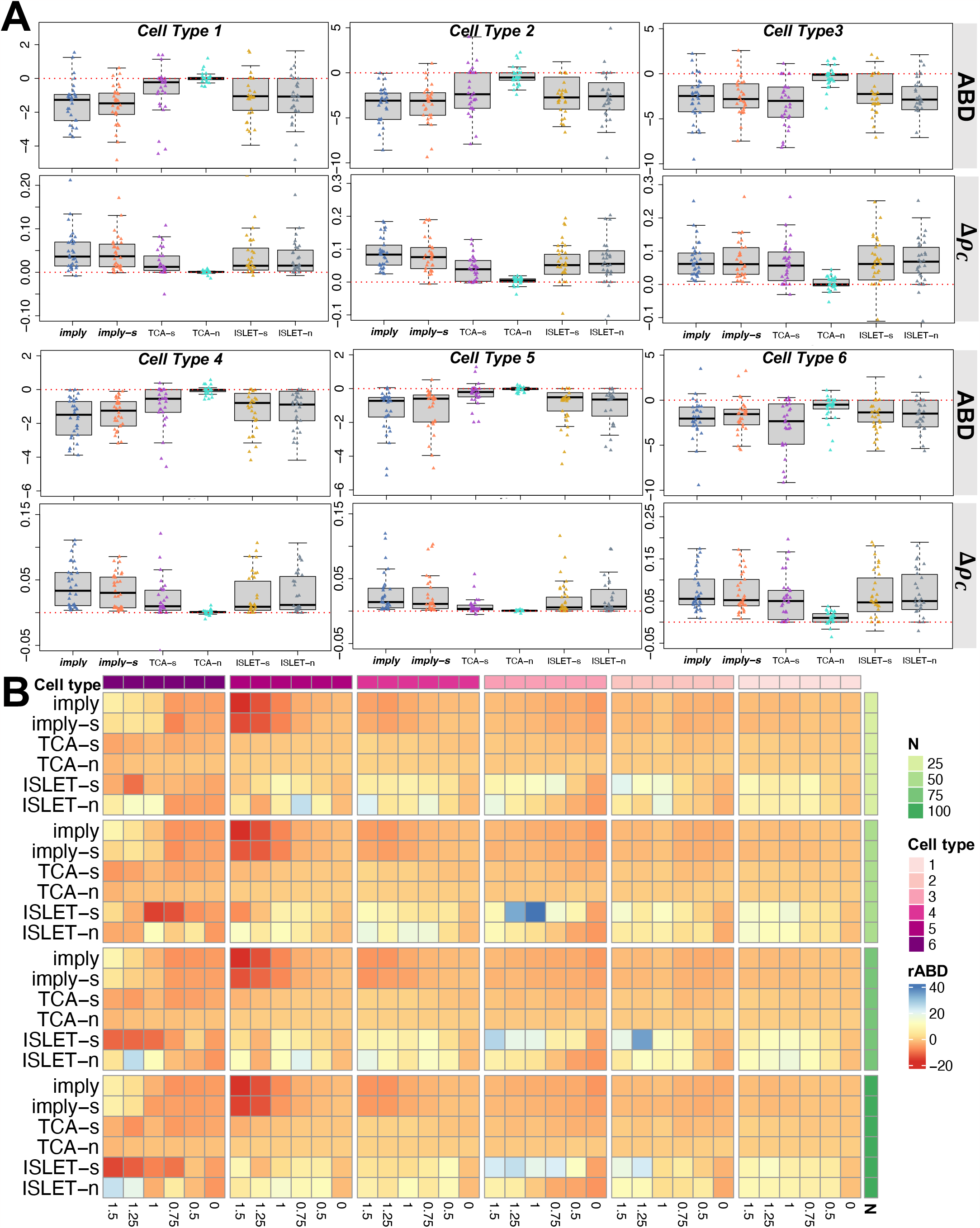
Cell-type resolution improvements in proportions estimated by ***imply*. (A)** Boxplots showing *ABD* (upper panel) and Δ*ρ*_C_ (lower panel), for cell types 1 to 6. **(B)** Heatmap showing the deconvolution improvement using the *rABD* metric, aggregated by cell types (top row) and sample sizes (right column), for various effect sizes (bottom row).

### Influential factors in deconvolution accuracy

We further zoom in to study how sample size, effect size, and subject-specific variation would affect personalized deconvolution. In **Figure 4A**, *ABD* and Δ*ρ*_C,E_ for ***imply***, together with ISLET-n and TCA-s, are presented across LFC ranging from 0 (null) to 1.5. ***imply*** consistently exhibits the lowest *ABD* in all scenarios and the highest Δ*ρ*_C,E_ in most settings. These results indicate the advantage of adopting personalized reference panels. In addition, ***imply*** provides the most stable (i.e., smallest variation) among the three methods as the effect size increases. **Figure 4B** shows the same metrics across various sample sizes. As expected, *ABD* decreases as the sample size increases. ***imply*** consistently maintains the highest Δ*ρ*_C,E_ across various sample sizes. In **Figure 4C**, we further investigate the Δ*ρ*_C,E_ alteration percentages, which are de-fined as 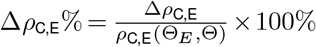 at different levels of SSV, which are annotated by the top row. We observe a robust pattern across different effect sizes, samples sizes, and SSVs, and conclude that ***imply*** and ***imply-s*** consistently provide the most outstanding concordance improvement.

**Figure 4.**
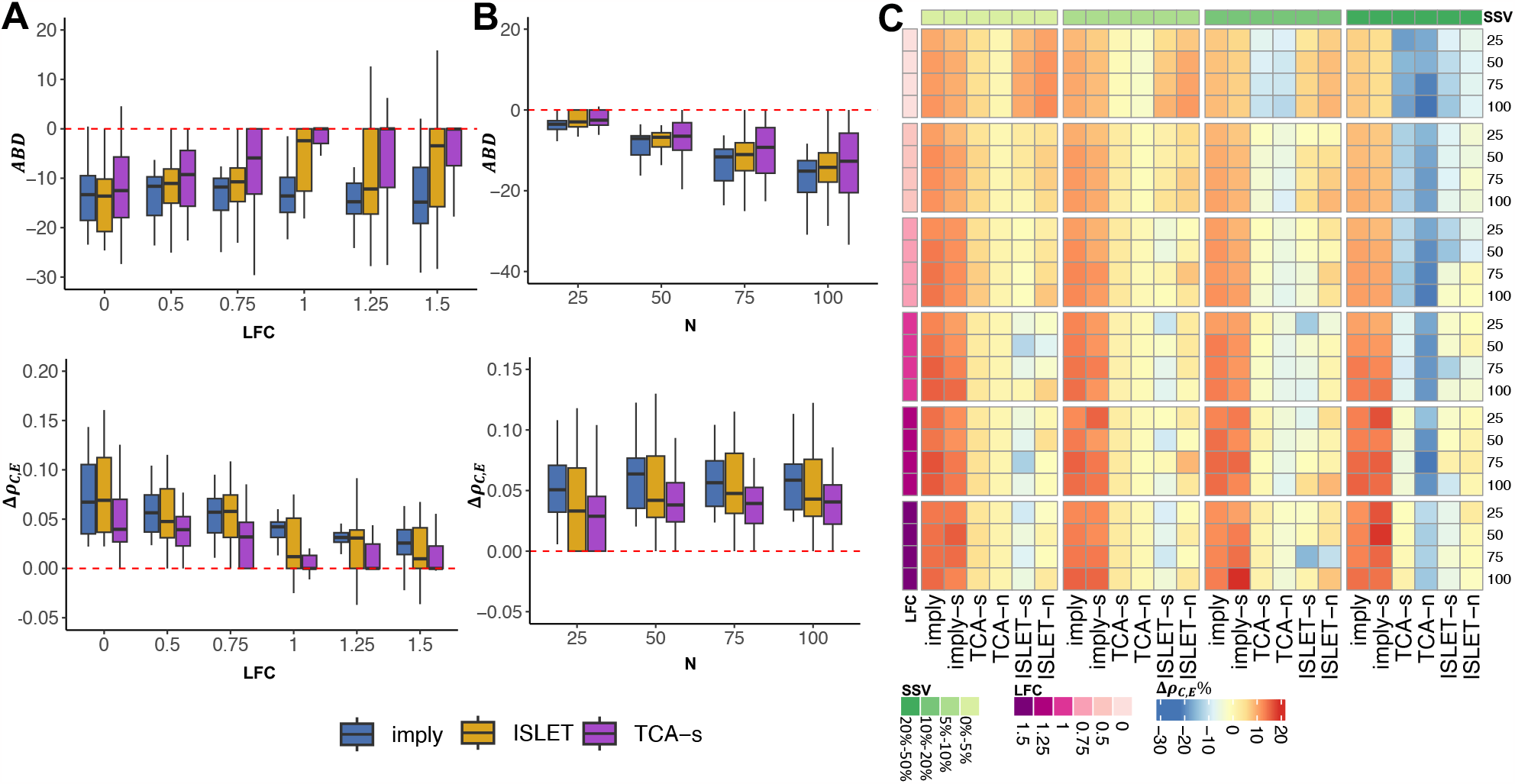
Effect size, sample size, and subject-specific variation affect deconvolution accuracy. **(A)** Boxplots of *ABD* (upper panel) and Δ*ρ*_C,E_ (lower panel) across three methods: ***imply***, ISLET-n, TCA-s, under different effect sizes (LFC): 0, 0.5, 0.75, 1, 1.25, and 1.5. **(B)** Similar to **(A)** but across various sample sizes per group: 25, 50, 75, 100. **(C)** Heatmap showing the relative Δ*ρ*_C,E_ across various combinations of sample sizes, effect sizes, and SSV. The color bars on the left and the top indicate the LFC and SSV, respectively. The number on the right indicates the sample size per group.

### Application of *imply* to longitudinal transcriptomic datasets

We applied ***imply*** to two consortia longitudinal transcriptomic datasets: one from Parkinson’s Disease Biomarker Program (PDBP) and the other one from The Environmental Determinants of Diabetes in the Young (TEDDY). The PDBP consortium has the repeatedly measured RNA-seq dataset, demographic and clinical information collected from patients with or without Parkinson’s Disease (PD) recruited from multiple medical centers and research institutions in the United States between November 2012 and August 2018. The PDBP cohort data were collected longitudinally over-time for each subject, allowing us to track changes in cell type composition and disease progression over time. In our study de-identified participants with at least three observations over time were retained. A total of 399 PD patients and 173 controls, with 2599 longitudinal samples over 2 years, were included. Longitudinal RNA samples in PDBP were extracted from the whole blood. Clinical data includes information about patients’ medical history, symptoms, disease status, total Montreal Cognitive Assessment (MoCA) scores, and MDS UPDRS part III motor scores. The TEDDY cohort is a multi-center pediatric study of Type 1 Diabetes (T1D). TEDDY cohort screened and enrolled participants with susceptibility of TID based on the Human Leukocyte Antigen (HLA) genotypes from six clinical centers in four countries (U.S., Finland, Germany, and Sweden). A total of 8,676 high-risk infants were enrolled from birth and followed every 3 months for blood sample collection and islet autoantibody (IAbs) measurement up to 4 years of age. Details of sample collection, RNA sequencing procedures, bioinformatics processing, and quality control in TEDDY are described in (Xhonneux et al., 2021). The longitudinal whole blood transcriptome data enable the ***imply*** deconvolution.

Figure 5. shows the deconvolution analysis results for PDBP and TEDDY real data. For PDBP dataset, the mean proportions across all visit times of six cell types, including B cell, Monocytes, CD4, CD8, NK cell, and other cells, are shown for cases and controls in **Figure 5A**. Here, B cell contributes the most among all six cell types, while NK contributes the least. The visualization suggests a higher CD8 proportions in the PD group than in the control group, while CD4 proportions in the PD groups are lower. **Figure 5B** displays the heatmap of Pearson correlations among the six cell types. B cells, monocytes, and CD4 all show negative pairwise correlations. **Figure 5C** shows box-plots of CD8 cell type proportions comparing case and control, at each time point. The median value of CD8 proportion in case is higher than that in control group at each time point. The CD4 and CD8 cell type pro-portions, broken down by the participant’s visit time of each subject, are shown in **Figure 5D** and **5E**, respectively. For CD4 cell type, the mean proportions in case group are lower than those in control group for each visit time. For CD8 cell type, the mean proportions among cases are higher than those among controls, for each visit time. These findings are well-aligned with previous studies where the PD patients showed elevated CD8 proportions and reduced CD4 proportions than controls (Wang et al., 2021; Baba et al., 2005; Galiano-Landeira et al., 2020). We also benchmarked ***imply*** with the existing method CIBERSORT as shown in **Figure 5F**. Using CIBERSORT, the *p*-value of the Wilcoxon Rank Sum test is 0.0111 and the median difference is −0.007 for CD8 proportions, between cases and controls. It incorrectly suggests that the CD8 cell type proportion of cases are lower than controls. In contrast, ***imply*** yields a *p*-value less than 10^−16^ and the median difference is 0.58, which shows the correct effect size direction. It also increases differential power between cases and controls, as shown in the ROC plot. We also explored the associations between the various cell type proportions and clinical outcomes, including total UPSIT score, total scores of Montreal Cognitive Assessment (MoCA), and MDS UPDRS part III motor scores, which provide additional assessments of patient’s cognitive and motor function in PD. Additionally, association studies with Cerebrospinal fluid (CSF) were conducted (results are in supplementary materials section 4.1). For the T1D study of TEDDY, the disease status (i.e., cases) of interest is the onset of pancreatic islet autoantibodies (IA). The longitudinal analysis of re-quantified cellular com-position identifies NK cell abundance as higher in males than females (*p <* 0.0001), as illustrated in **Figure 5H**. Previous research in TEDDY reported a higher risk of IA being associated with viral infection during the first 6 months of life (Vehik et al., 2019). The sex difference in NK cell fraction in **Figure 5H** could be a consequence of early-life vaccination or viral infection (Cheng et al., 2023), since infants are exposed to exogenous antigens and have a high susceptibility to infections. In this analysis, we use longitudinal samples of IA cases and controls collected at the age of 9-21 months, and compare deconvoluted cell fractions between groups by linear mixed effect model. **Figure 5I** shows that the NK cell proportions are significantly lower (*p <* 0.0001) in the participants who developed IA at a young age compared to controls, while this trend is not observed in the initial cell abundance estimated by CIBERSORT (*p* = 0.77, supplementary material section 4.2). The relative higher NK cell abundance in males (vs. females) and controls (vs. cases) among TEDDY participants is consistent with the previous finding that males have a lower risk of autoimmunity than females (Markle et al., 2013).

Furthermore, we perform a downstream csDE genes analysis on IA status based on the ***imply*** -deconvoluted cell type fraction, using ISLET (Feng et al., 2023) with FDR*<* 0.1. The cell type proportions improved by ***imply*** enabled the detection of DE genes in CD4 T cells and identified more NK-cell-specific DE genes (*n >* 300) compared to a previous csDE genes testing result (*n* = 30) based on the proportions deconvoluted by Auto-GeneS (Aliee and Theis, 2021). The IA-csDE genes based on the improved cell fractions include the markers for multiple T cell receptors (e.g., TRBV, TRDV, TRGV, TRJV) and the genes regulating immune responses such as *CAMP* and *CRK*. The *CAMP* gene expression was found to be associated with serum levels of vitamin D in the studies of innate immunity (Lowry et al., 2020; Gombart et al., 2009; de Oliveira et al., 2022), while the TEDDY cohort also reported a strong linkage between vitamin D and the risk of IA (Li et al., 2021). Protein *CRK* is involved in NK cells inhibitory receptor signaling and modulates the signaling of activating receptors, which may function as a two-way molecular switch to control NK cell-mediated cytotoxicity (Nabekura et al., 2018; Liu, 2014).

**Figure 5.**
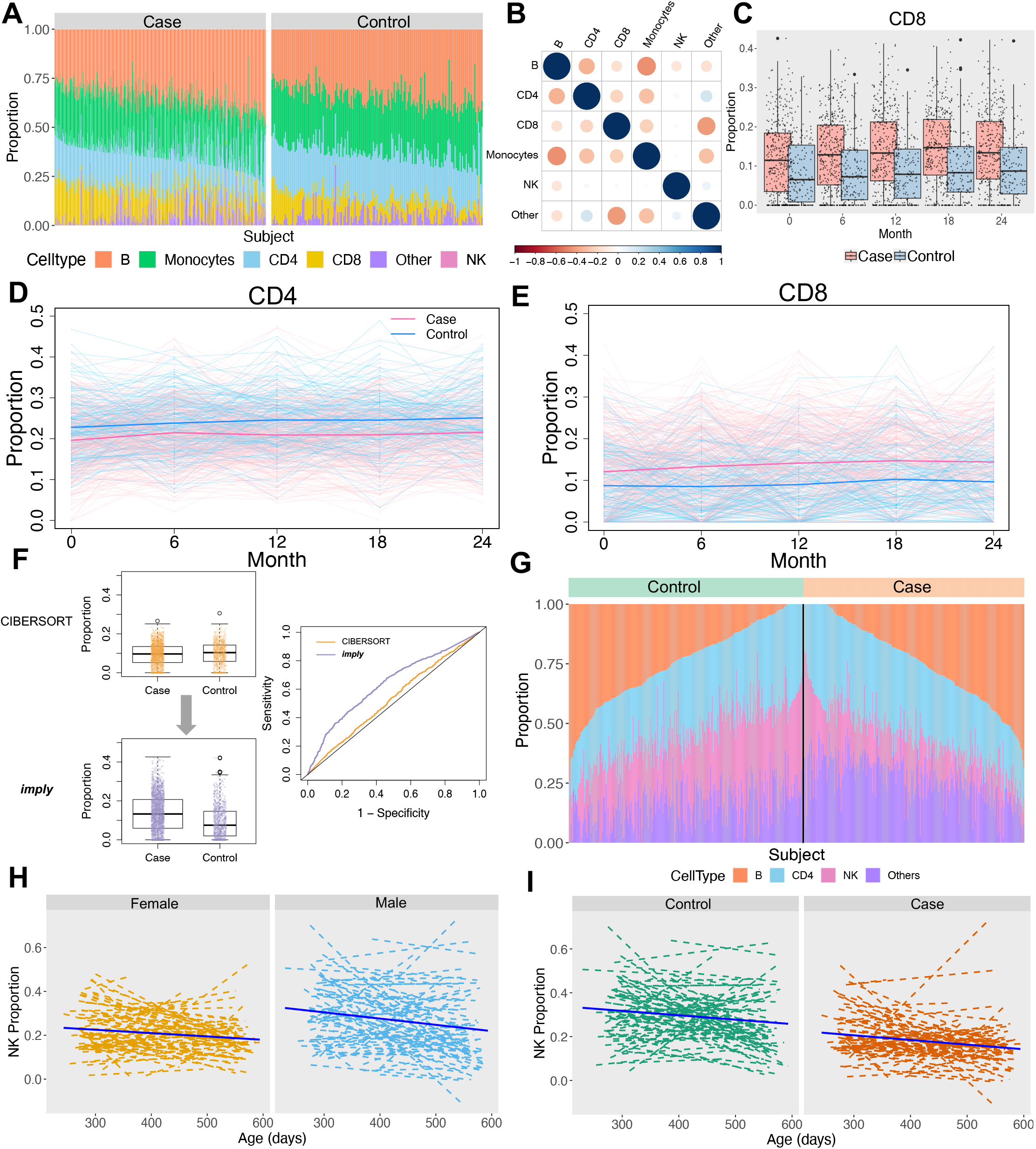
Phenotype-associated cell type disparities from PDBP and TEDDY consortia. PDBP: Parkinson’s Disease Biomarker Program. TEDDY: The Environmental Determinants of Diabetes in the Young. **(A)**: Cell type proportions for all subjects, separated by Parkinson’s Disease (PD) status in PDBP dataset. The bar represents the mean cell type proportions across all visit times, for each subject. **(B)**: Pearson correlations of cell type proportions between six cell types, among all individuals. **(C)**: Distribution comparisons of CD8 cell type proportions between PD cases and controls, at each visit time. **(D)**: Deconvoluted CD4 cell type proportions along with study participants’ visit time. PD cases (pink) and controls (blue) are illustrated by both individual background lines (thin) and foreground lines (thick). **(E)**: Same as in **(D)** but for CD8 cell type proportions. **(F)**: Grand comparison of CD8 cell type proportions between PD cases and healthy controls, using CIBERSORT and our method ***imply*** . Results from ***imply*** show a larger effect size, more significant test statistics, and increased discriminative capacity. **(G)**: Cell type proportions for all subjects, separated by pancreatic islet autoantibodies (IA) status, in TEDDY dataset. The bar represents the mean cell type proportions across all visit times, for each subject. **(H)**: NK cell proportions along infant’s age (in days) at sample collection, for female and male subjects. Average fitted lines (solid) overlay individual-specific lines (dashed). **(I)**: Same as in **(H)** but separated by IA case and control status.

## DISCUSSION

In this work, we present our statistical framework ***imply*** to conduct cell-type deconvolution in bulk data using personalized panels. Our method ***imply*** leverages the repeated bulk RNA-seq samples to purify personalized reference transcriptome, and then jointly quantifies the cell abundances across multiple samples per individual. We show the advantage of using personalized reference panels by extensively *in silico* simulation studies and the analytical results of two large-scale lon-gitudinal consortia. ***imply*** can produce more accurate and realistic deconvolution results.

The computational deconvolution of admixed bulk tissue samples is drawing substantial interests in *-omics*. The interest is growing as deconvolution methodology are being developed, and as increasingly large datasets are becoming available with and without repeated measures. We are among the first to consider personalized reference panels in deconvolution. Our computational framework optimizes the usages of shared information in longitudinal samples from each subject. Alternative machine learning approaches, such as Expectation-Maximization (EM) and non-negative matrix factorization algorithms, could also extract personalized reference panels and have been implemented in ISLET (Feng et al., 2023) and CIBERSORTx (Newman et al., 2019). Nevertheless, these methods lack the conciseness and computational efficiency exhibited by the proposed linear mixed-effects modeling framework.

A limitation of ***imply*** is the requirement of an initial signature matrix as the input in *Stage I*, which could affect the initial cell type abundance estimation as the input for downstream. An alternative approach is to initialize cell fractions by external multi-subject reference cell count data, such as single-cell profiling and labeling, flow cytometry, or imaging. For some genes, the random effect variance estimation may shrink towards zero, likely due to the adoption of penalized MLE. For such scenarios, the cell-type-specific heterogeneity between individuals would not be fully recovered. Furthermore, the intra-individual heterogeneity was not considered in reference panel recovery. This is because our present work was motivated by the bulk transcriptome of longitudinal blood samples, many of which were collected from healthy controls. In those scenarios, the underlying pure gene expression panel for each subject is relatively stable over time. Our previous work (Feng et al., 2023) suggests that the intra-individual cell-type-specific heterogeneity, when assessing using longitudinal PBMC scRNA-seq data, is trivial when compared with inter-individual variation. Hence, our future work will include the curation of longitudinal scRNA-seq data from distinct tissue types or disease populations and the incorporation of potential variations between time points at cell type resolution.

## DATA AVAILABILITY

- ***imply*** is implemented and integrated into a R/Bioconductor package ISLET, which is available at https://bioconductor.org/packages/ISLET.
- The PDBP bulk transcriptome and related clinical data are publicly available on request to AMP-PD at https://amp-pd.org
- The TEDDY bulk transcriptome dataset has been deposited in NCBI’s database of Genotypes and Phenotypes (dbGaP) with the primary accession code phs001442.v3.p2

## ACKNOWLEDGEMENTS

The TEDDY study is funded by the National Institute of Diabetes and Digestive and Kidney Diseases, National Institute of Allergy and Infectious Diseases, National Institute of Child Health and Human Development, National Institute of Environmental Health Sciences, Centers for Disease Control and Prevention, and JDRF. We thank the TEDDY study data coordinating center at Health Informatics Institute, University of South Florida, for data processing and sharing. The PDBP consortium is supported by the National Institute of Neurological Disorders and Stroke (NINDS) at the National Institutes of Health. A full list of PDBP investigators can be found at https://pdbp.ninds.nih.gov/policy. The PDBP investigators have not participated in reviewing the data analysis or content of the manuscript. Data used in the preparation of this article were obtained from the Accelerating Medicine Partnership (AMP) Parkinson’s Disease (AMP PD) Knowledge Platform.

