## Supplementary Materials for "*imply*: improving cell-type deconvolution accuracy using personalized reference profiles"

Guanqun Meng, Yue Pan, Wen Tang, Lijun Zhang, Ying Cui,
Fredrick R. Schumacher, Ming Wang, Rui Wang, Sijia He, Jeffrey Krischer,
Qian Li*, and Hao Feng*

### 1 Method Details

#### 1.1 $\nu$-SVR Objective Function Constrains

The $\nu$-SVR has been shown to work well in well-received deconvolution algorithms Newman et al. [2015, 2019] and the proportion of support vectors could be manually tuned by $\epsilon$ rooted into the objective function (main Figure 1). The support vectors are those observations outside the $\epsilon$-tube (main Figure 1), which determine the hyperplane boundaries fitting the maximum number of observations Schölkopf et al. [2000]. Also, observations within the $\epsilon$-tube receive no penalty in the loss function. The objective function (in main text) is subject to the constraints, $\forall j$:

$$\begin{aligned} \boldsymbol{y}_{\cdot ni} - f(\boldsymbol{\theta}_{E,ni\cdot}) \leq \epsilon + \xi_j \\ f(\boldsymbol{\theta}_{E,ni\cdot}) - \boldsymbol{y}_{\cdot ni} \leq \epsilon + \xi_j^* \end{aligned}$$

Here, $\epsilon$ and $C$ are tunable margin and positive constant which help to prevent overfitting by controlling the penalties caused by observations outside the $\epsilon$-tube (Figure 1). $\xi_j$ and $\xi_j^*$ are slack variables denoting deviance from the margin and introducing extra error spaces to satisfy the potential infeasible but mandatory constraints. The concept is like a soft-margin concept in support vector machine (SVM) classification. The loss function ignores the prediction errors within $\epsilon$-tube and is defined:

$$L_\epsilon = \begin{cases} 0 & , \text{if } |(\boldsymbol{y}_{\cdot ni})'_j - f(\boldsymbol{\theta}_{E,ni\cdot})_j| < \epsilon \\ |(\boldsymbol{y}_{\cdot ni})'_j - f(\boldsymbol{\theta}_{E,ni\cdot})_j| - \epsilon & , \text{otherwise} \end{cases}$$

#### 1.2 Mixed Effect Model Design Matrices

Here $\boldsymbol{X}$ and $\boldsymbol{A}$ are design matrices, each has dimension $T \times 2K$ and $T \times NK$, respectively. These matrices are employed in the linear mixed-effect regression to model the fixed-effect $\boldsymbol{\beta}_g$ and the random-effect $\boldsymbol{u}_g$. Additionally, $\boldsymbol{a}_{nk} = (\hat{\theta}_{E,n1k}, \hat{\theta}_{E,n2k}, \ldots \hat{\theta}_{E,nt_nk})'$ denotes the initial cell type abundance information.

$$
\boldsymbol{X} =
\begin{bmatrix}
\hat{\theta}_{E,111} & \hat{\theta}_{E,112} & \dots & \hat{\theta}_{E,11K} & z_1\hat{\theta}_{E,111} & z_1\hat{\theta}_{E,112} & \dots & z_1\hat{\theta}_{E,11K} \\
\hat{\theta}_{E,121} & \hat{\theta}_{E,122} & \dots & \hat{\theta}_{E,12K} & z_1\hat{\theta}_{E,121} & z_1\hat{\theta}_{E,112} & \dots & z_1\hat{\theta}_{E,12K} \\
\vdots & & \ddots & & & & & \\
\hat{\theta}_{E,1t_11} & \hat{\theta}_{E,1t_12} & \dots & \hat{\theta}_{E,1t_1K} & z_1\hat{\theta}_{E,1t_11} & z_1\hat{\theta}_{E,1t_12} & \dots & z_1\hat{\theta}_{E,1t_1K} \\
\vdots & & & \ddots & & & & \\
\hat{\theta}_{E,N11} & \hat{\theta}_{E,N12} & \dots & \hat{\theta}_{E,N1K} & z_N\hat{\theta}_{E,N11} & z_1\hat{\theta}_{E,N12} & \dots & z_1\hat{\theta}_{E,N1K} \\
\hat{\theta}_{E,N21} & \hat{\theta}_{E,N22} & \dots & \hat{\theta}_{E,N2K} & z_N\hat{\theta}_{E,N21} & z_1\hat{\theta}_{E,N22} & \dots & z_1\hat{\theta}_{E,N2K} \\
\vdots & & & & \ddots & & & \\
\hat{\theta}_{E,Nt_N1} & \hat{\theta}_{E,Nt_N2} & \dots & \hat{\theta}_{E,Nt_NK} & z_N\hat{\theta}_{E,Nt_N1} & z_N\hat{\theta}_{E,Nt_N2} & \dots & z_1\hat{\theta}_{E,Nt_NK}
\end{bmatrix}
$$

$$
\boldsymbol{A} =
\begin{bmatrix}
\boldsymbol{a}_{11} & \mathbf{0} & \mathbf{0} & \dots & \mathbf{0} & \boldsymbol{a}_{12} & \mathbf{0} & \mathbf{0} & \dots & \mathbf{0} & \boldsymbol{a}_{1K} & \mathbf{0} & \mathbf{0} & \dots & \mathbf{0} \\
\mathbf{0} & \boldsymbol{a}_{21} & \mathbf{0} & \dots & \mathbf{0} & \mathbf{0} & \boldsymbol{a}_{22} & \mathbf{0} & \dots & \mathbf{0} & \mathbf{0} & \boldsymbol{a}_{2K} & \mathbf{0} & \dots & \mathbf{0} \\
\vdots & \ddots & & & \vdots & & \ddots & & & & \vdots & \ddots & & & \vdots \\
\vdots & & & \ddots & \vdots & & & & \ddots & & \vdots & & & \ddots & \vdots \\
\mathbf{0} & \mathbf{0} & \mathbf{0} & \dots & \boldsymbol{a}_{N1} & \mathbf{0} & \mathbf{0} & \mathbf{0} & \dots & \boldsymbol{a}_{N2} & \mathbf{0} & \mathbf{0} & \mathbf{0} & \dots & \boldsymbol{a}_{NK}
\end{bmatrix}
$$

#### 1.3 Evaluation Metric

Lin's concordance correlation coefficient was designed for univariate outcome but not specifically for high-dimension compositional measurements, where the relative abundances within a subject could be dependent and subject to the positiveness and unit-sum constraints. To address these aspects, Cui et al. Cui et al. [2021] propose a general formulation of CCC by substituting the expected squared difference to Euclidean distance (shown in the main text) or Aitchison distance Aitchison et al. [2000], which is a popular distance measure for paired compositional data, defined below:

$$
d_A^2(\boldsymbol{\Theta}, \hat{\boldsymbol{\Theta}}) = \sum_{k=1}^{K} \left[ \log\left( \frac{\hat{\boldsymbol{\Theta}}^{(k)}}{h(\hat{\boldsymbol{\Theta}})} \right) - \log\left( \frac{\boldsymbol{\Theta}^{(k)}}{h(\boldsymbol{\Theta})} \right) \right]^2
$$

where $h(\boldsymbol{\Theta}) = \left(\prod_{l=1}^{K} \boldsymbol{\Theta}^{(l)}\right)^{\frac{1}{K}}$ indicates the geometric mean of the elements of $\boldsymbol{\Theta}$. If the proposed method leads to increased concordance, we can expect to observe positive differences in Lin's CCC and its adapted versions when comparing the improved cell proportion estimation with the initial estimation.

### 2 Simulation Details

#### 2.1 Real Data Selection and Cell-Specific Underlying Parameters' Estimation

The gene expression specific to each cell type is obtained from an actual study (GSE60424), which investigates differences in the transcriptome among six immune cell types (Neutrophils, Monocytes, B-cells, CD4 T cells, CD8 T cells, and Natural Killer cells) between healthy individuals and patients with immune-associated diseases (specifically, Type 1 Diabetes, Amyotrophic Lateral Sclerosis, Sepsis, and Multiple Sclerosis patients) as reported in Linsley et al. [2014]. The underlying expression parameters for a given cell type $k$ and gene $g$ are estimated using the *estParam* function from the *PROPER package* Wu et al. [2015]. This estimation assumes that the RNA-seq read counts follow a negative binomial distribution ($NB(\mu_{gk}, \phi_{gk})$), where $\mu_{gk}$ and $\phi_{gk}$ represent the mean expression and biological dispersion for gene $g$, respectively.

The initial parameters specific to cells and genes are initially refined using Human Genome-Wide annotations. Genes lacking corresponding genetic symbols from org.HS.eg.db Carlson [2021] are excluded, resulting in 30,080 remaining genes. Subsequently, these gene-specific underlying estimators are further filtered based on specific thresholds ($\hat{\mu}_{gk} > 1$ and $\hat{\phi}_{gk} > 0.001$), resulting in a total of 1,540 underlying gene estimations that serve as references for simulations. Tables 1 and 2 present the empirical mean and covariance matrix for the cell-specific filtered gene expression parameters.

| | B-Cell | CD4 | CD8 | Monocytes | Neutrophils | NK |
|---|---|---|---|---|---|---|
| $\bar{\boldsymbol{\mu}}_m$ | 4.53 | 4.48 | 4.56 | 4.72 | 4.49 | 4.53 |
| $\bar{\boldsymbol{\mu}}_\phi$ | -4.04 | -4.06 | -3.60 | -4.00 | -3.26 | -3.82 |

Table 1: The averages for underlying gene expression parameters of the 1,540 genes that remain for each cell type in healthy samples.

| $\hat{\boldsymbol{\Sigma}}_m$ ($\hat{\boldsymbol{\Sigma}}_\phi$) | B-Cell | CD4 | CD8 | Monocytes | Neutrophils | NK |
|---|---|---|---|---|---|---|
| B-Cell | 2.19 (2.00) | 1.73 (0.95) | 1.69 (0.75) | 1.50 (0.61) | 1.30 (0.55) | 1.47 (0.71) |
| CD4 | | 2.09 (2.02) | 1.98 (1.02) | 1.38 (0.63) | 1.26 (0.60) | 1.63 (0.78) |
| CD8 | | | 2.02 (1.90) | 1.42 (0.64) | 1.25 (0.57) | 1.73 (0.80) |
| Monocytes | | | | 2.16 (1.72) | 2.03 (0.74) | 1.41 (0.57) |
| Neutrophils | | | | | 4.05 (2.32) | 1.44 (0.56) |
| NK | | | | | | 1.80 (2.01) |

Table 2: Variances and covariances among the underlying gene expression parameters of the 1,540 retained genes across various cell types in healthy samples.

Celltype1-celltype6 will mask cell type names for further simulations. *Multivariate Gaussian* distributions are used to simulate two matrices representing cell-specific gene expression parameters for each subject. The underlying cell-specific gene expression matrix is generated through a re-parameterized *Gamma Distribution* for each sample, as depicted in the equations below.

$$
\begin{aligned}
&\boldsymbol{M} \sim MVN(\bar{\boldsymbol{\mu}}_m, \hat{\boldsymbol{\Sigma}}_m) \\
&\boldsymbol{\Phi} \sim MVN(\bar{\boldsymbol{\mu}}_\phi, \hat{\boldsymbol{\Sigma}}_\phi) \\
&\boldsymbol{P} \sim \Gamma(\frac{1}{\exp(\boldsymbol{\Phi})}, \exp(\boldsymbol{M}) \times \exp(\boldsymbol{\Phi}))
\end{aligned}
$$

#### 2.2 Cell mixture proportion generation

Cell proportion estimations rely on 16 different single-cell RNA sequencing labeling studies, each with varying numbers of cell types ranging from 3 to 13. Using bootstrapping and resampling techniques, we created a pooled sample of 80 cells, allowing us to empirically derive cell-type compositions from this cell-label pool. This procedure was repeated 200 times, resulting in a 200×6 cell composition matrix. The matrix was solved using the *Dirichlet Distribution* via the *dirichlet.mle* function from the *sirt* package Robitzsch [2022]. This yielded a vector of cell proportion distributional parameters for control samples, denoted as $\boldsymbol{\alpha}_{ctrl} = [8.85, 6.49, 5.98, 5.28, 4.22, 3.85]$. For the case samples, the *Dirichlet* parameters were generated by redistributing the sum of $\alpha$ values ($\alpha_{Total} = \sum_{k=1}^{6} \alpha_{ctrl}$). The cell proportion distributional parameters for cases are: $\boldsymbol{\alpha}_{case} = [1.90, 2.25, 2.10, 5.72, 7.33, 15.37]$.

### 3 Simulation Results

Additional simulation results are presented here in the supplementary section, including six evaluation metrics: absolute bias difference ($ABD$, Section 4.1), relative absolute bias difference ($rABD$%, Section 4.2), correlation difference ($CD$, Section 4.3), differences of Lin's CCC ($\Delta\rho_{\text{C}}$, Section 4.4), differences of Euclidean-based CCC ($\Delta\rho_{\text{C,E}}$, Section 4.5), and differences of Aitchison-based CCC ($\Delta\rho_{\text{C,A}}$, Section 4.6) across different scenarios. In these scenarios, subject-specific variation ($SSV$) is set at levels of 0-5% and 10-20%. The effect sizes and sample sizes per group are respectively reflected by: $LFC = 0$, 0.5, 1.5, and $N = 25$, 75, 100. Each plot shows a particular sample size with various combinations of LFC and SSV.

#### 3.1 Absolute Bias Difference (ABD)

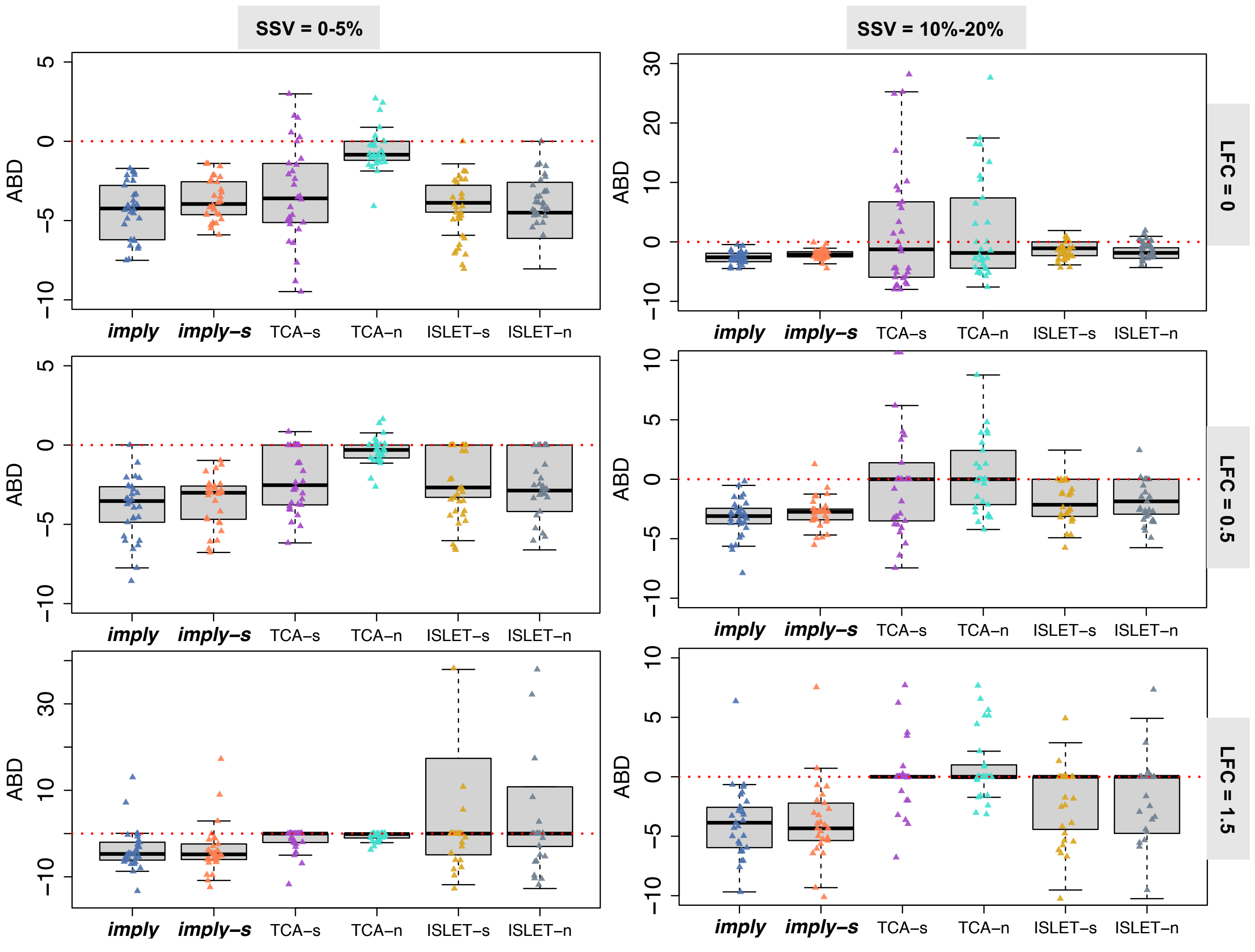

Figure 1: The absolute bias difference ($ABD$) is examined across various levels of Subject-Specific Variations ($SSV$s) represented in columns and effect sizes depicted in rows, focusing specifically on a sample size ($N$) of 25.

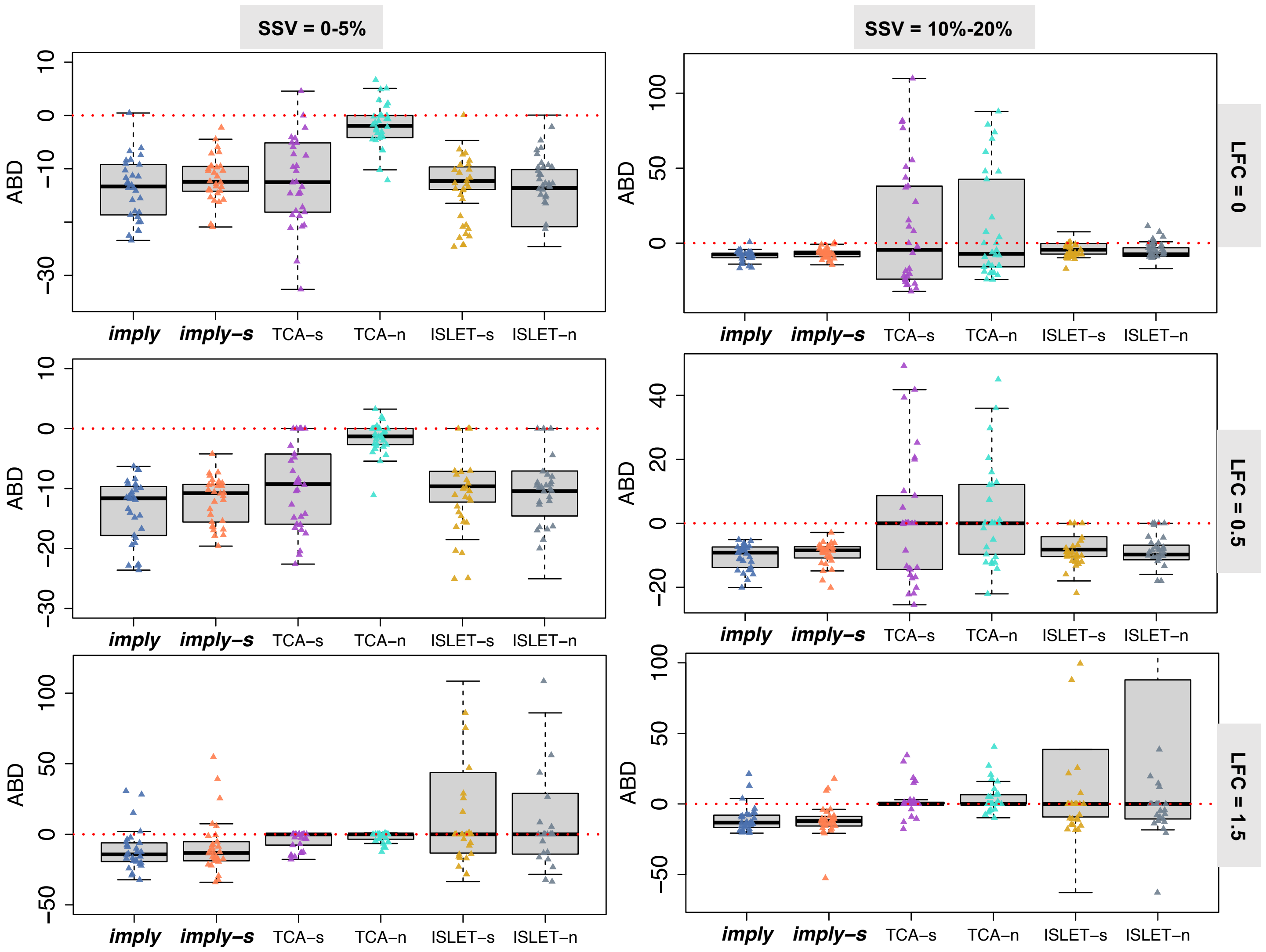

Figure 2: The absolute bias difference ($ABD$) is examined across various levels of Subject-Specific Variations ($SSV$s) represented in columns and effect sizes depicted in rows, focusing specifically on a sample size ($N$) of 75.

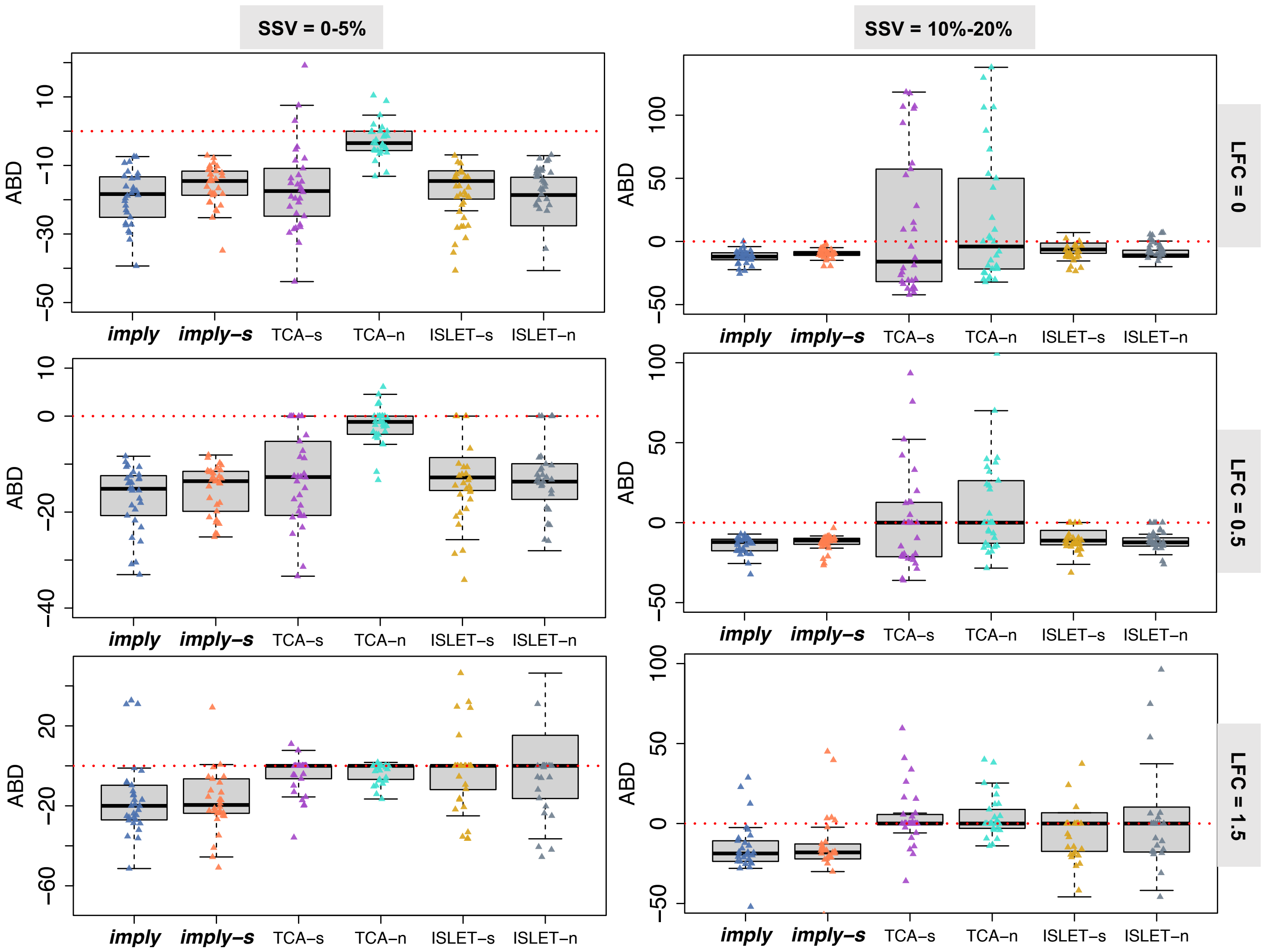

Figure 3: The absolute bias difference ($ABD$) is examined across various levels of Subject-Specific Variations ($SSV$s) represented in columns and effect sizes depicted in rows, focusing specifically on a sample size ($N$) of 100.

#### 3.2 Relative absolute bias difference (rABD%)

For both $ABD$ and $rABD\%$, if they are less than zero, it signifies that the evaluated method decrease the estimation bias. Furthermore, lower values additionally suggest better models' performances.

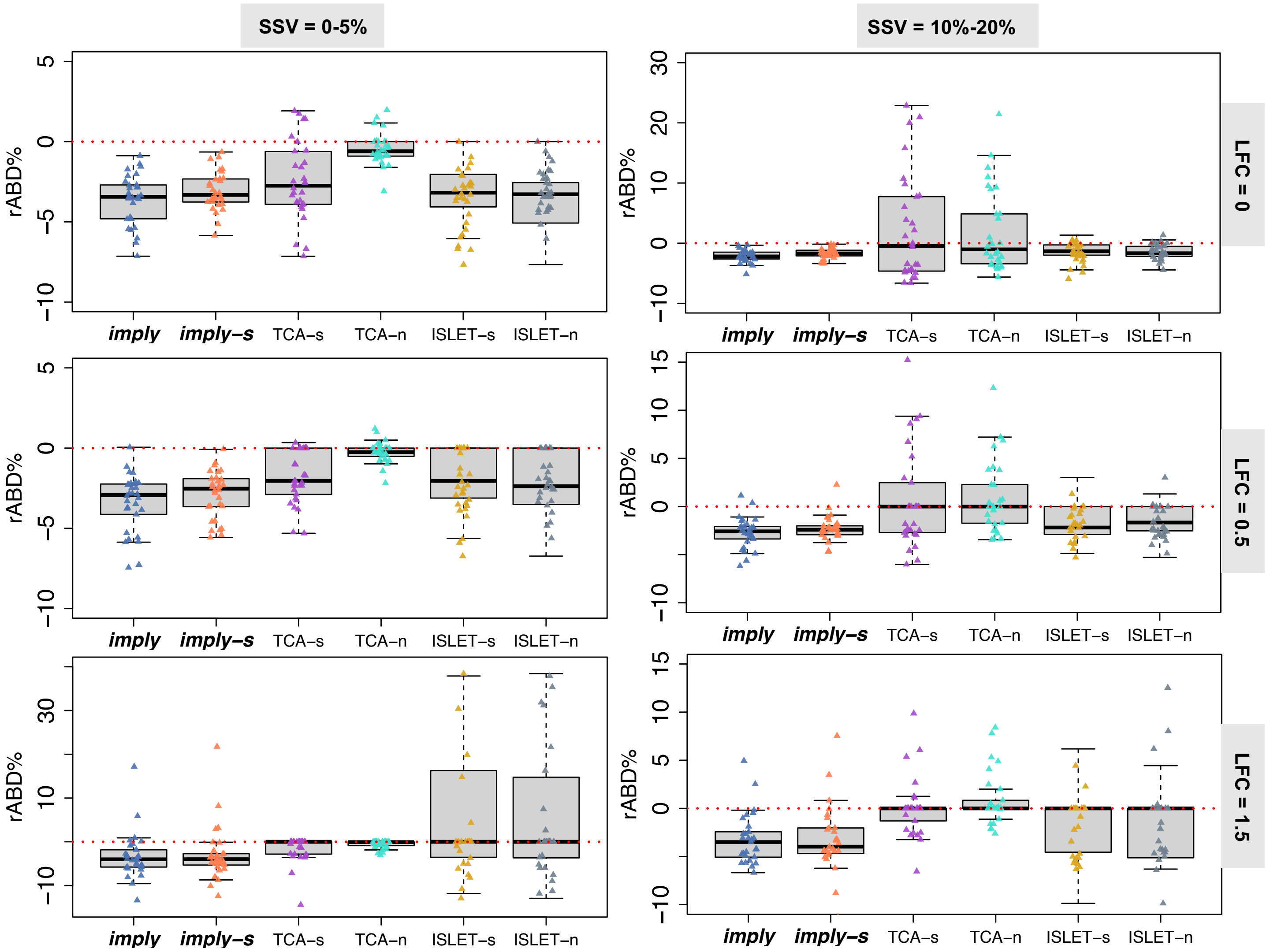

Figure 4: The relative absolute bias difference ($rABD\%$) is examined across various levels of Subject-Specific Variations ($SSV$s) represented in columns and effect sizes depicted in rows, focusing specifically on a sample size ($N$) of 25.

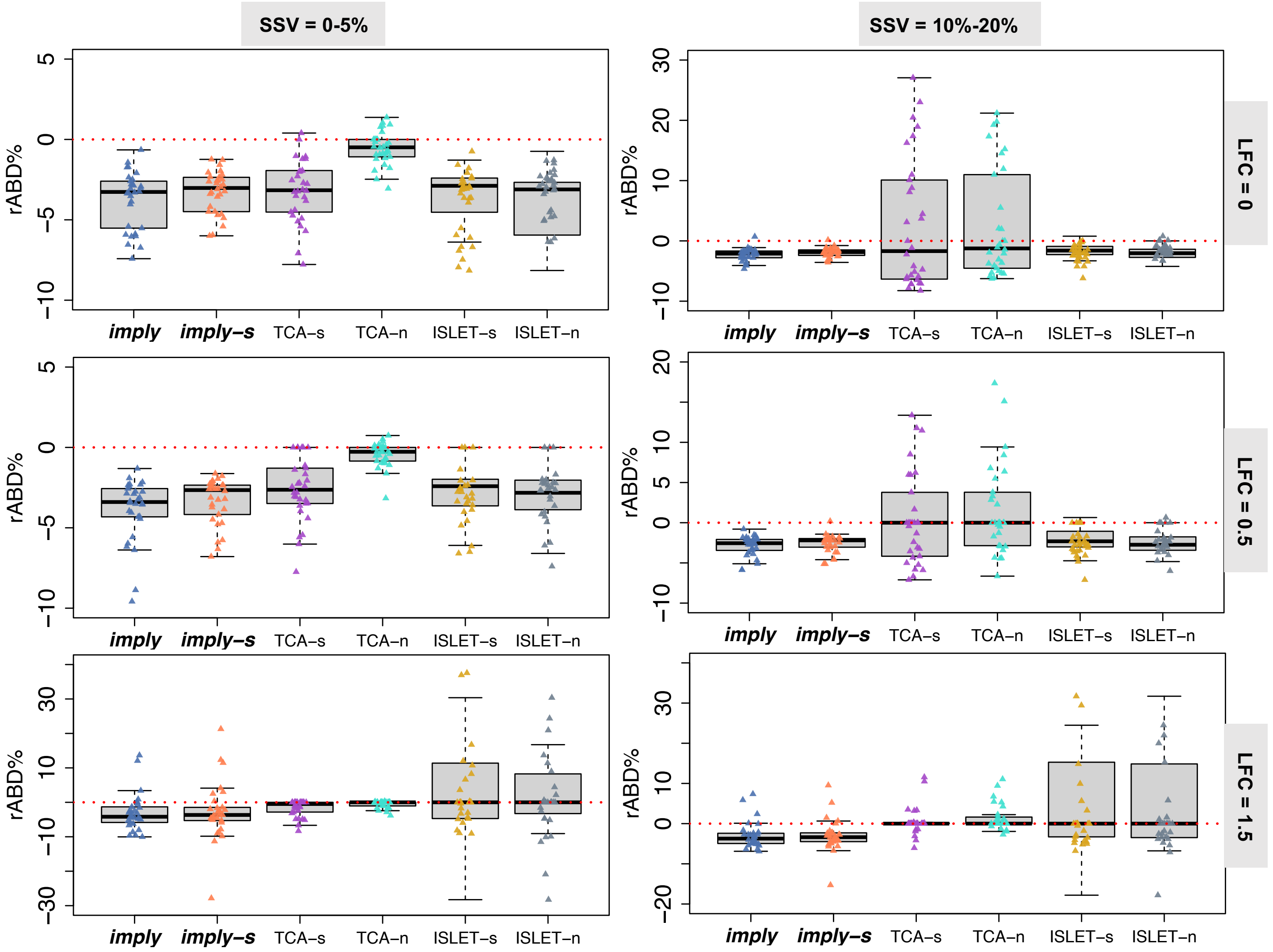

Figure 5: The relative absolute bias difference ($rABD\%$) is examined across various levels of Subject-Specific Variations ($SSVs$) represented in columns and effect sizes depicted in rows, focusing specifically on a sample size ($N$) of 75.

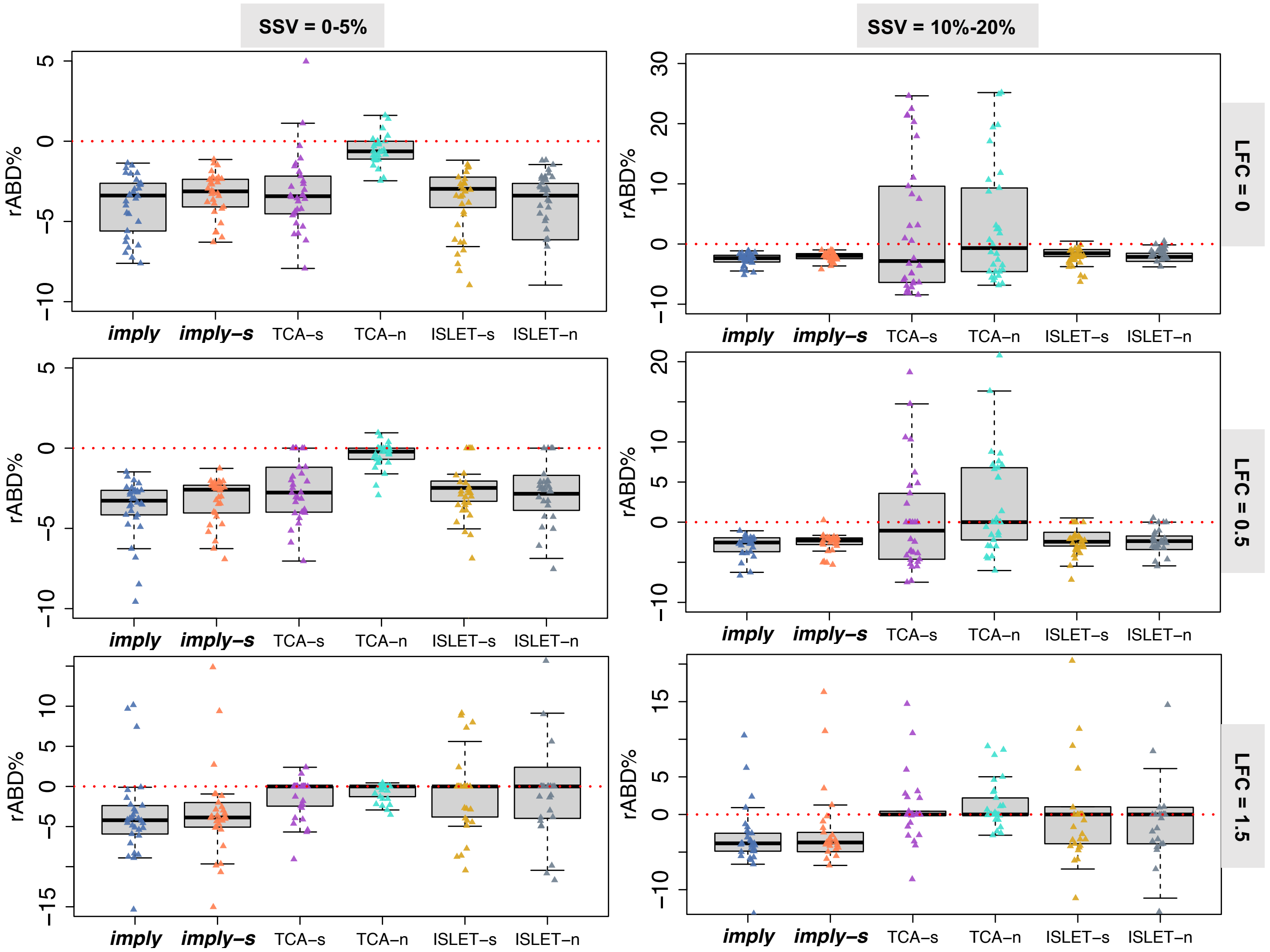

Figure 6: The relative absolute bias difference ($rABD\%$) is examined across various levels of Subject-Specific Variations ($SSVs$) represented in columns and effect sizes depicted in rows, focusing specifically on a sample size ($N$) of 100.

##### 3.3 Correlation Differences

When $CD > 0$, it implies that the evaluated method enhances the correlation between estimated cell type proportions and the ground truth. A greater value indicates a more promising performance.

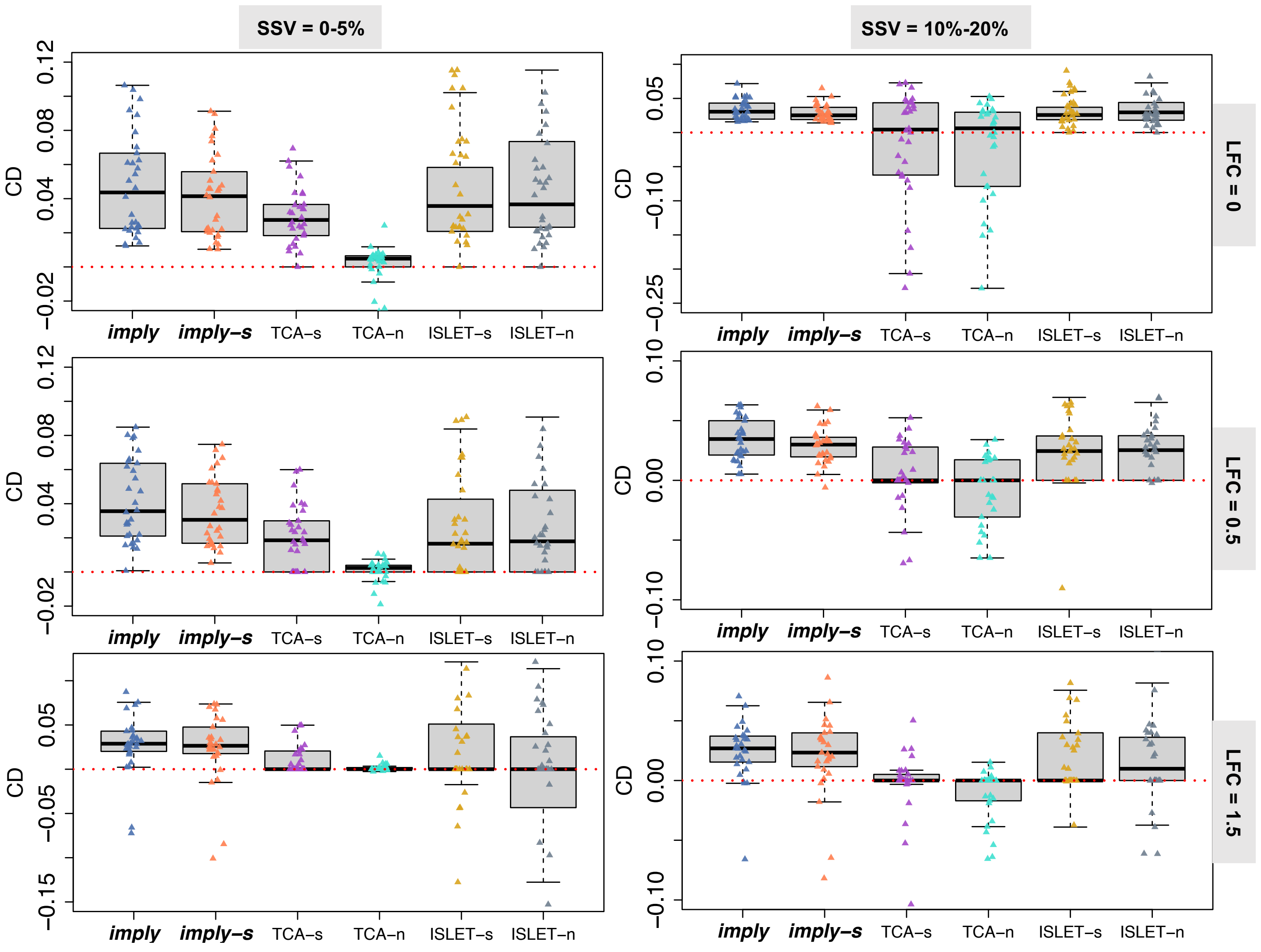

Figure 7: The correlation difference ($CD$) is examined across various levels of Subject-Specific Variations ($SSV$s) represented in columns and effect sizes depicted in rows, focusing specifically on a sample size ($N$) of 25.

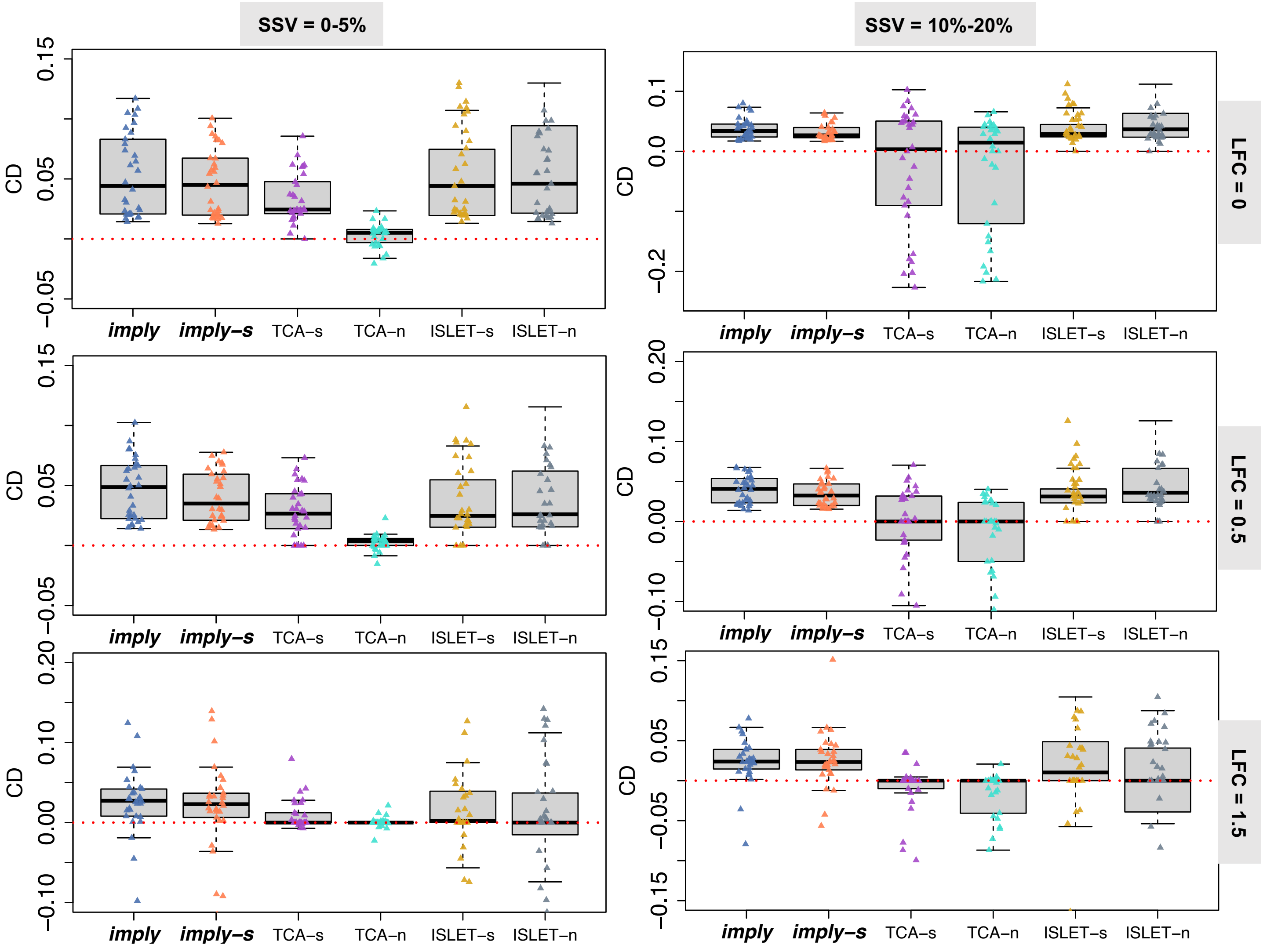

Figure 8: The correlation difference ($CD$) is examined across various levels of Subject-Specific Variations ($SSV$s) represented in columns and effect sizes depicted in rows, focusing specifically on a sample size ($N$) of 75.

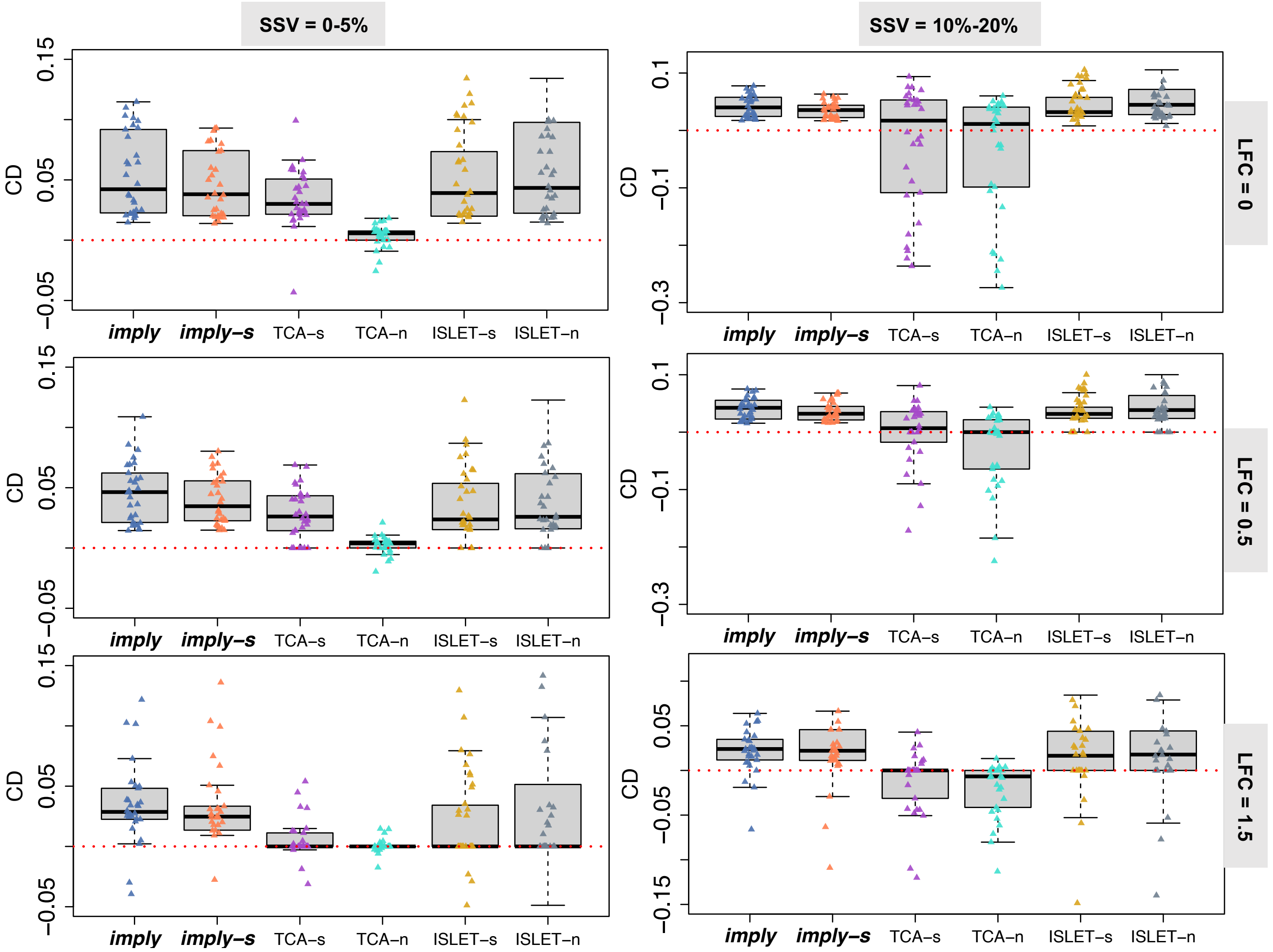

Figure 9: The correlation difference ($CD$) is examined across various levels of Subject-Specific Variations ($SSV$s) represented in columns and effect sizes depicted in rows, focusing specifically on a sample size ($N$) of 100.

##### 3.4 Lin's CCC

If the evaluated method improve estimation concordance, we would anticipate observing positive values in $\Delta\rho_C$, $\Delta\rho_{C,E}$ and $\Delta\rho_{C,A}$.

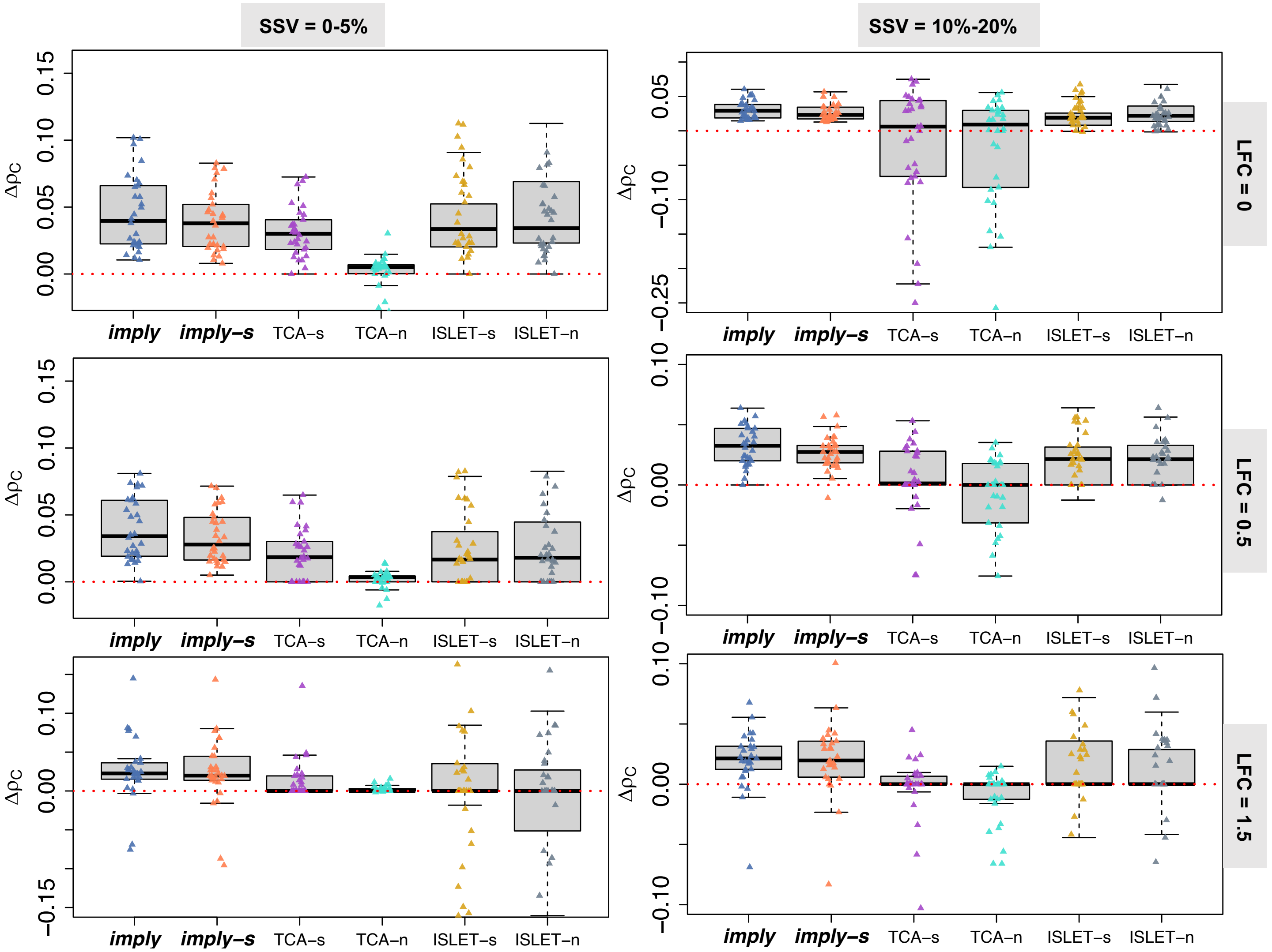

Figure 10: The difference of Euclidean-based CCC ($\Delta\rho_{C,E}$) is examined across various levels of Subject-Specific Variations ($SSV$s) represented in columns and effect sizes depicted in rows, focusing specifically on a sample size ($N$) of 25.

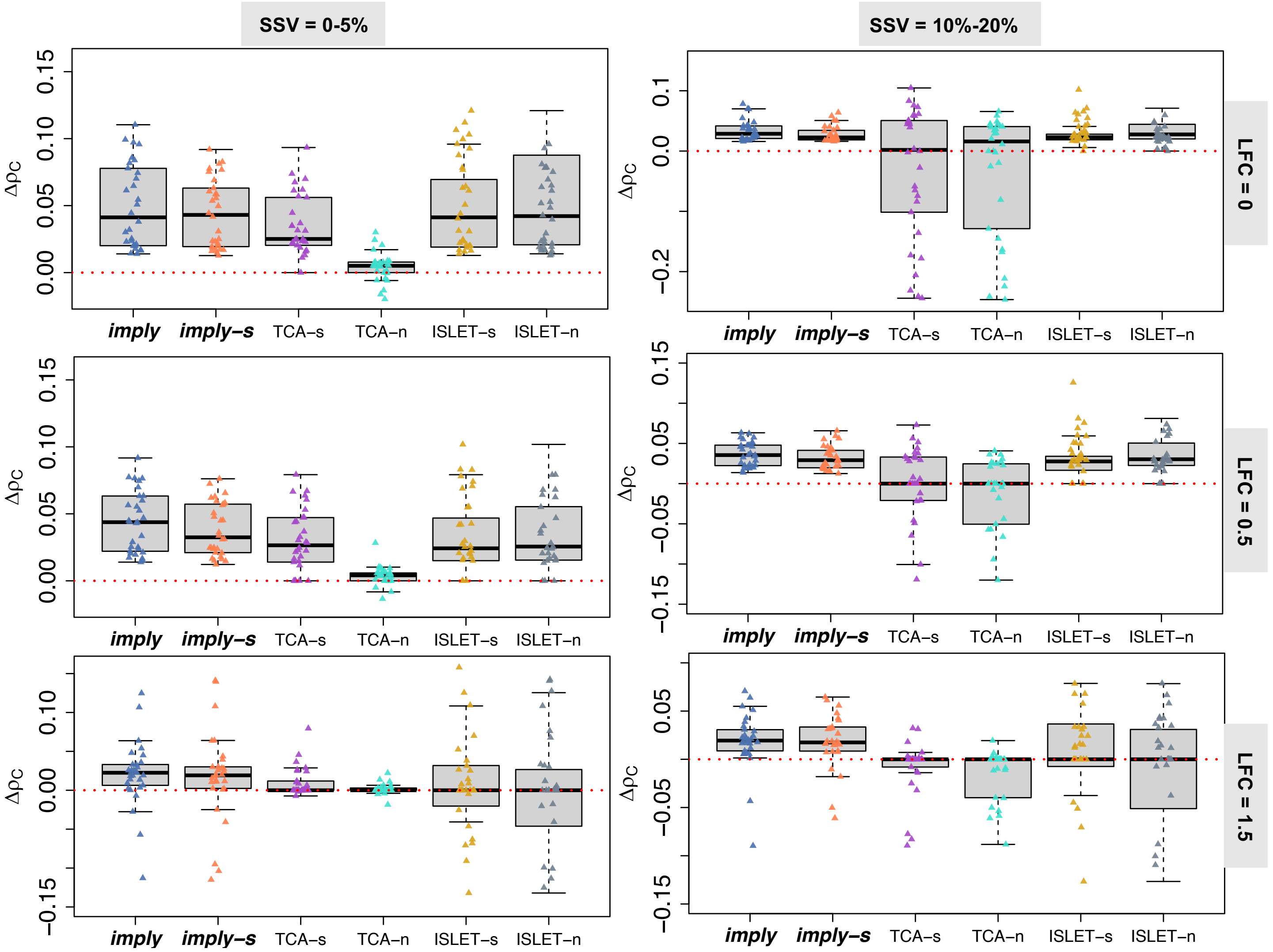

Figure 11: The difference of Euclidean-based CCC ($\Delta\rho_{\mathrm{C,E}}$) is examined across various levels of Subject-Specific Variations ($SSV$s) represented in columns and effect sizes depicted in rows, focusing specifically on a sample size ($N$) of 75.

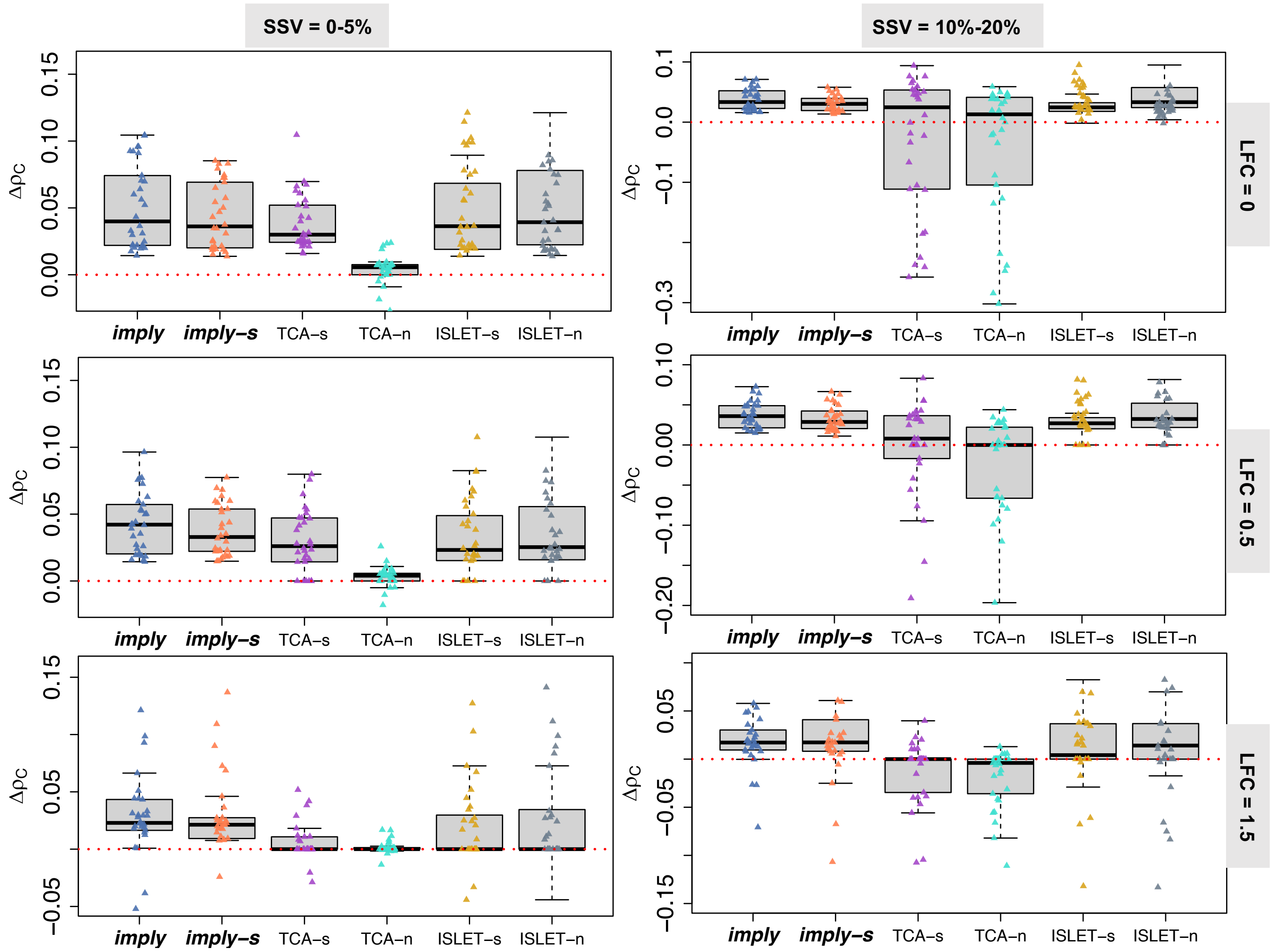

Figure 12: The difference of Euclidean-based CCC ($\Delta\rho_{\mathrm{C,E}}$) is examined across various levels of Subject-Specific Variations ($SSV$s) represented in columns and effect sizes depicted in rows, focusing specifically on a sample size ($N$) of 100.

##### 3.5 Euclidean Based-CCC

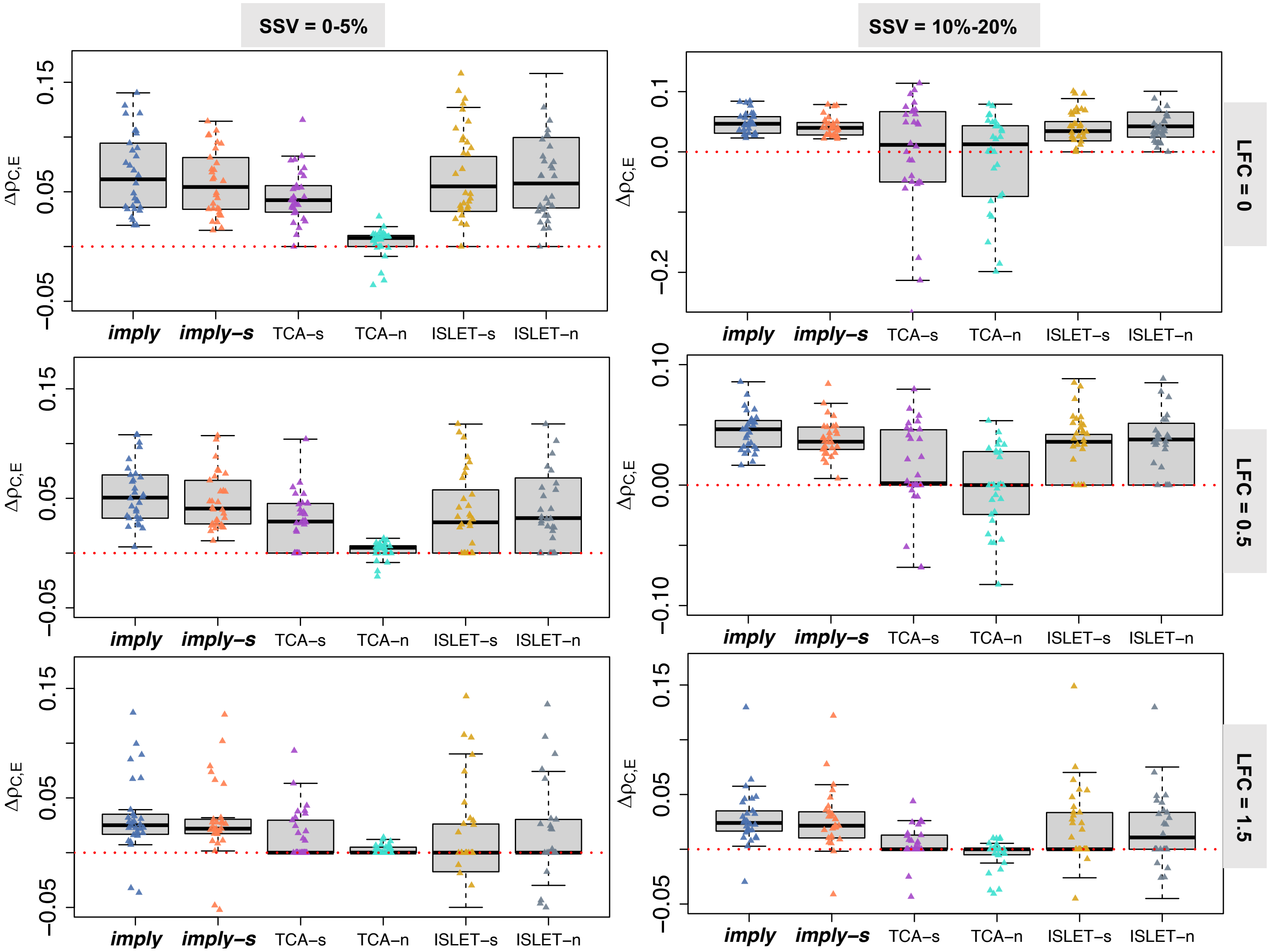

Figure 13: The difference of Euclidean-based CCC ($\Delta\rho_{\mathrm{C,E}}$) is examined across various levels of Subject-Specific Variations ($SSV$s) represented in columns and effect sizes depicted in rows, focusing specifically on a sample size ($N$) of 25.

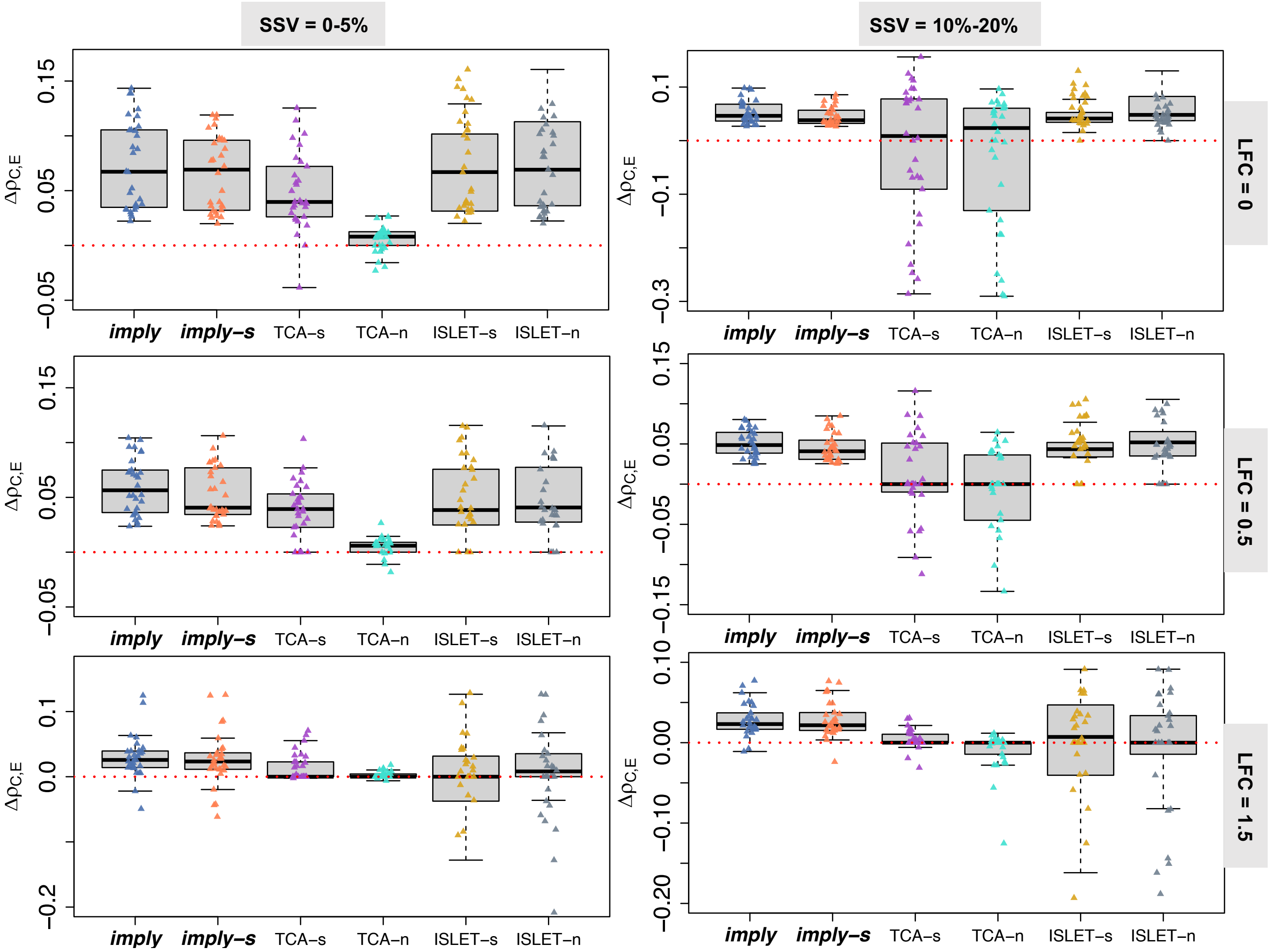

Figure 14: The difference of Euclidean-based CCC ($\Delta\rho_{\mathrm{C,E}}$) is examined across various levels of Subject-Specific Variations ($SSV$s) represented in columns and effect sizes depicted in rows, focusing specifically on a sample size ($N$) of 75.

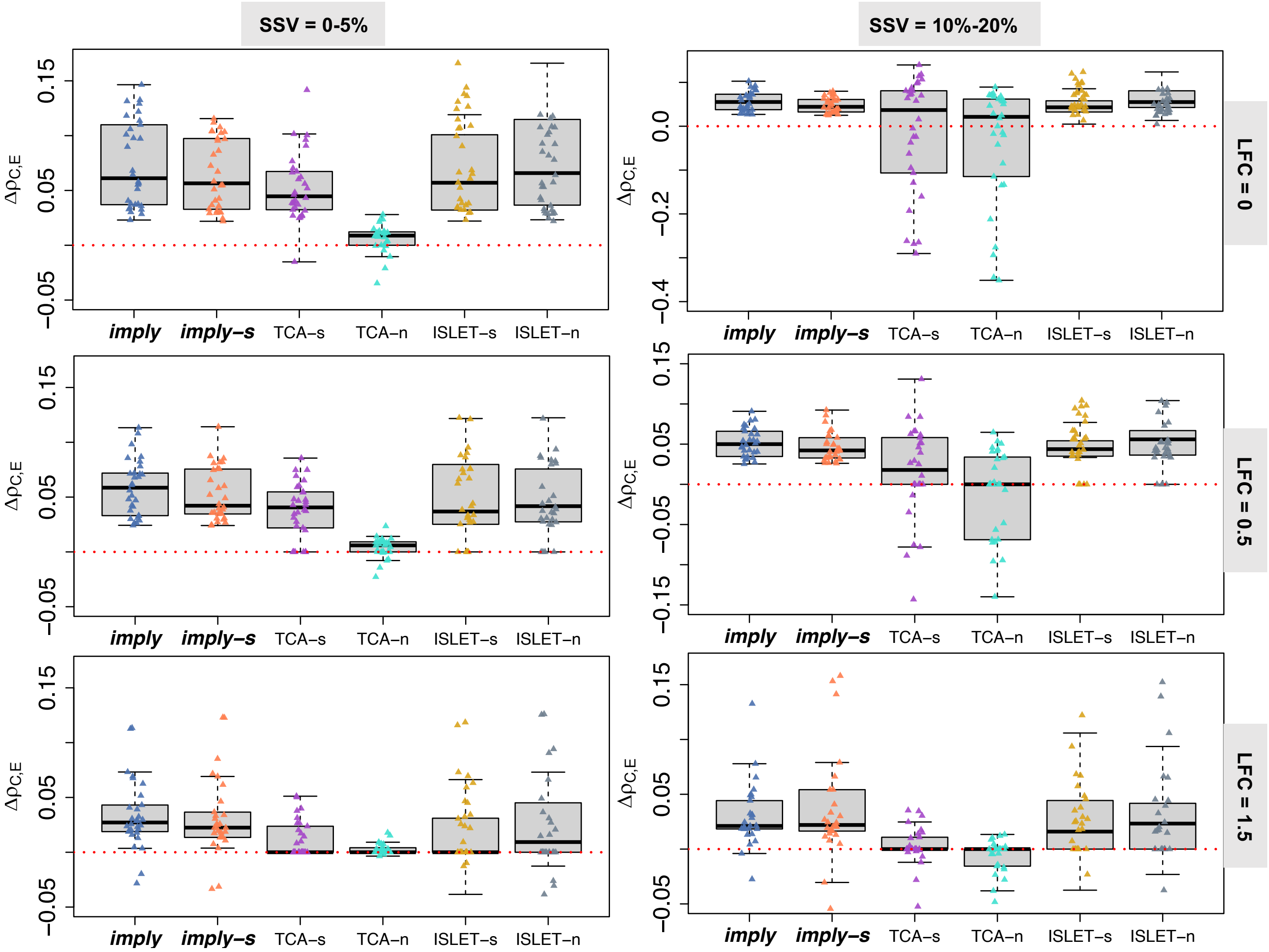

Figure 15: The difference of Euclidean-based CCC ($\Delta\rho_{\mathrm{C,E}}$) is examined across various levels of Subject-Specific Variations ($SSV$s) represented in columns and effect sizes depicted in rows, focusing specifically on a sample size ($N$) of 100.

##### 3.6 Aitchison Based-CCC

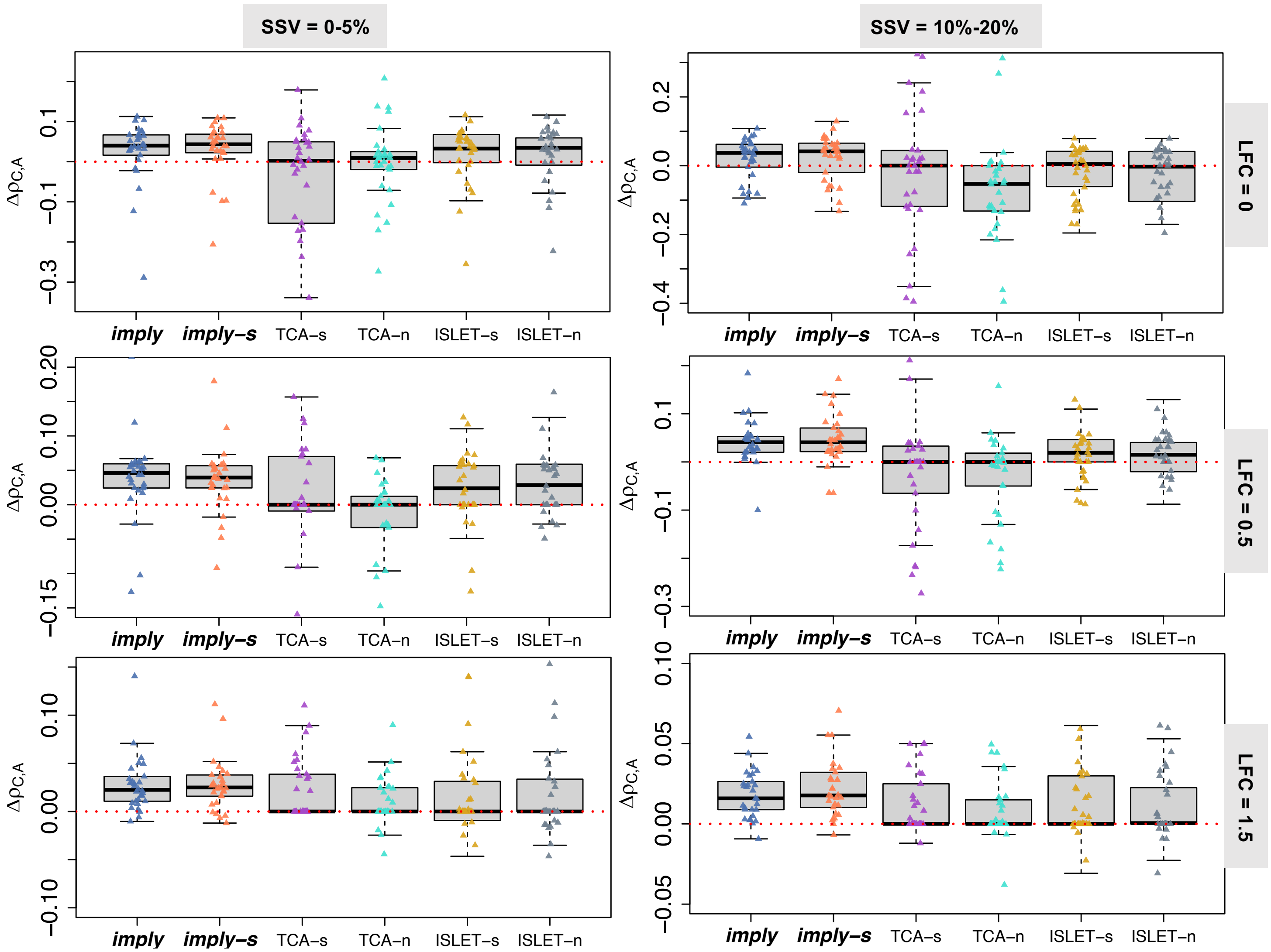

Figure 16: The difference of Aitchison-based CCC ($\Delta\rho_{\mathrm{C,A}}$) is examined across various levels of Subject-Specific Variations ($SSV$s) represented in columns and effect sizes depicted in rows, focusing specifically on a sample size ($N$) of 25.

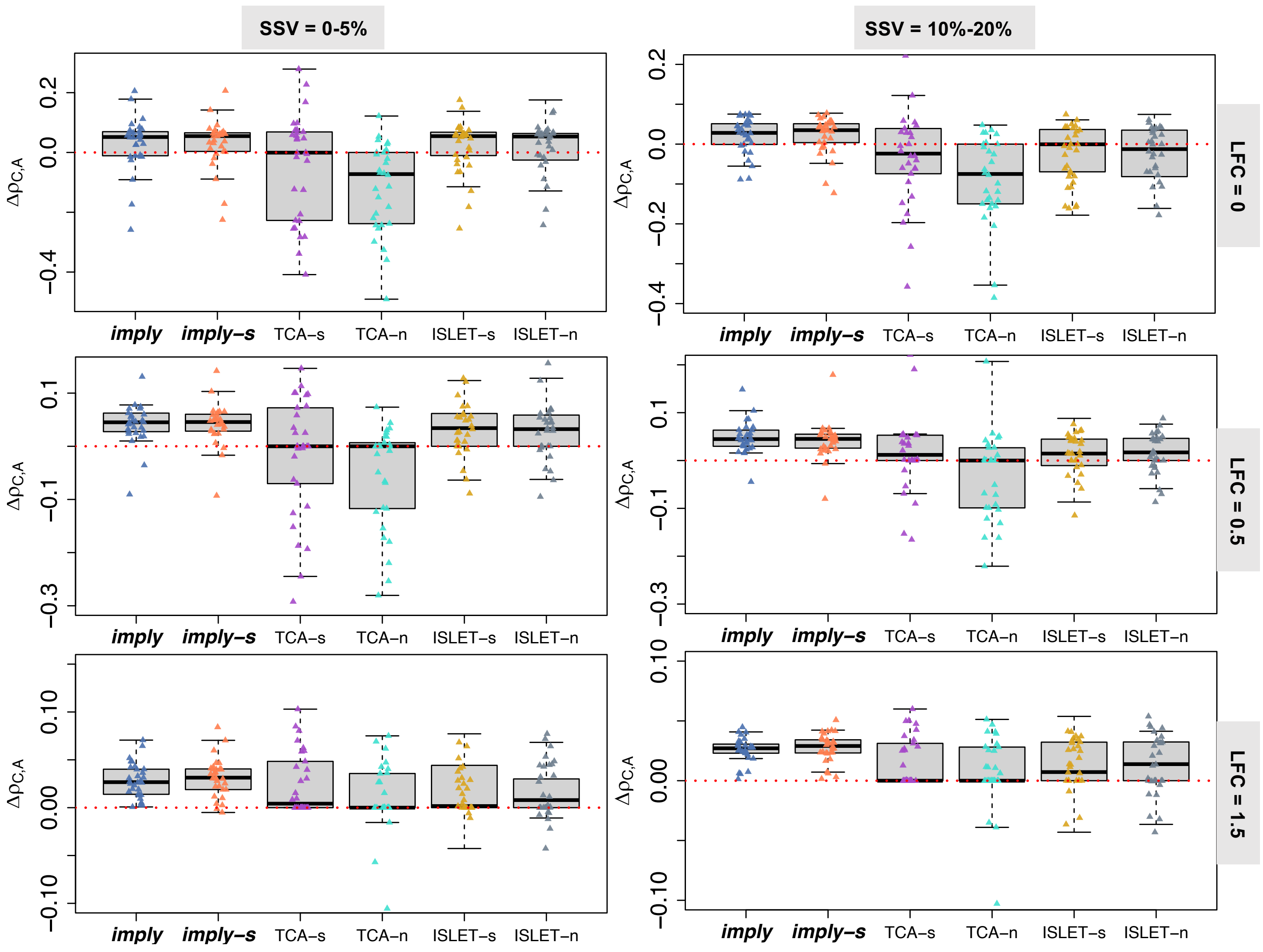

Figure 17: The difference of Aitchison-based CCC ($\Delta\rho_{\mathrm{C,A}}$) is examined across various levels of Subject-Specific Variations ($SSV$s) represented in columns and effect sizes depicted in rows, focusing specifically on a sample size ($N$) of 75.

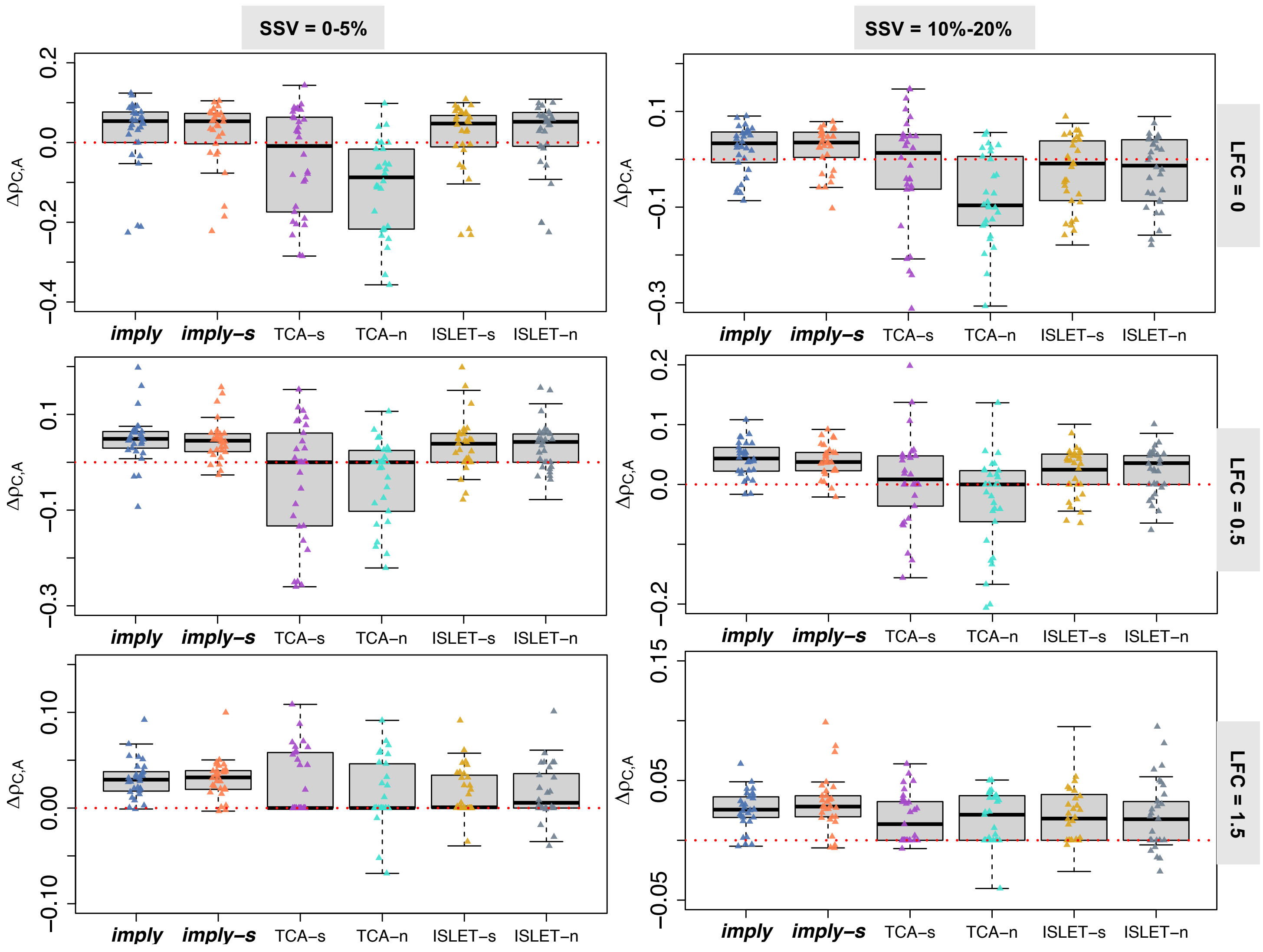

Figure 18: The difference of Aitchison-based CCC ($\Delta\rho_{\mathrm{C,A}}$) is examined across various levels of Subject-Specific Variations ($SSV$s) represented in columns and effect sizes depicted in rows, focusing specifically on a sample size ($N$) of 100.

### 4 Real Data Analysis

#### 4.1 PDBP: Parkinson's Disease Biomarker Program

Here, we investigate the correlation between different cell type proportions and clinical indicators.

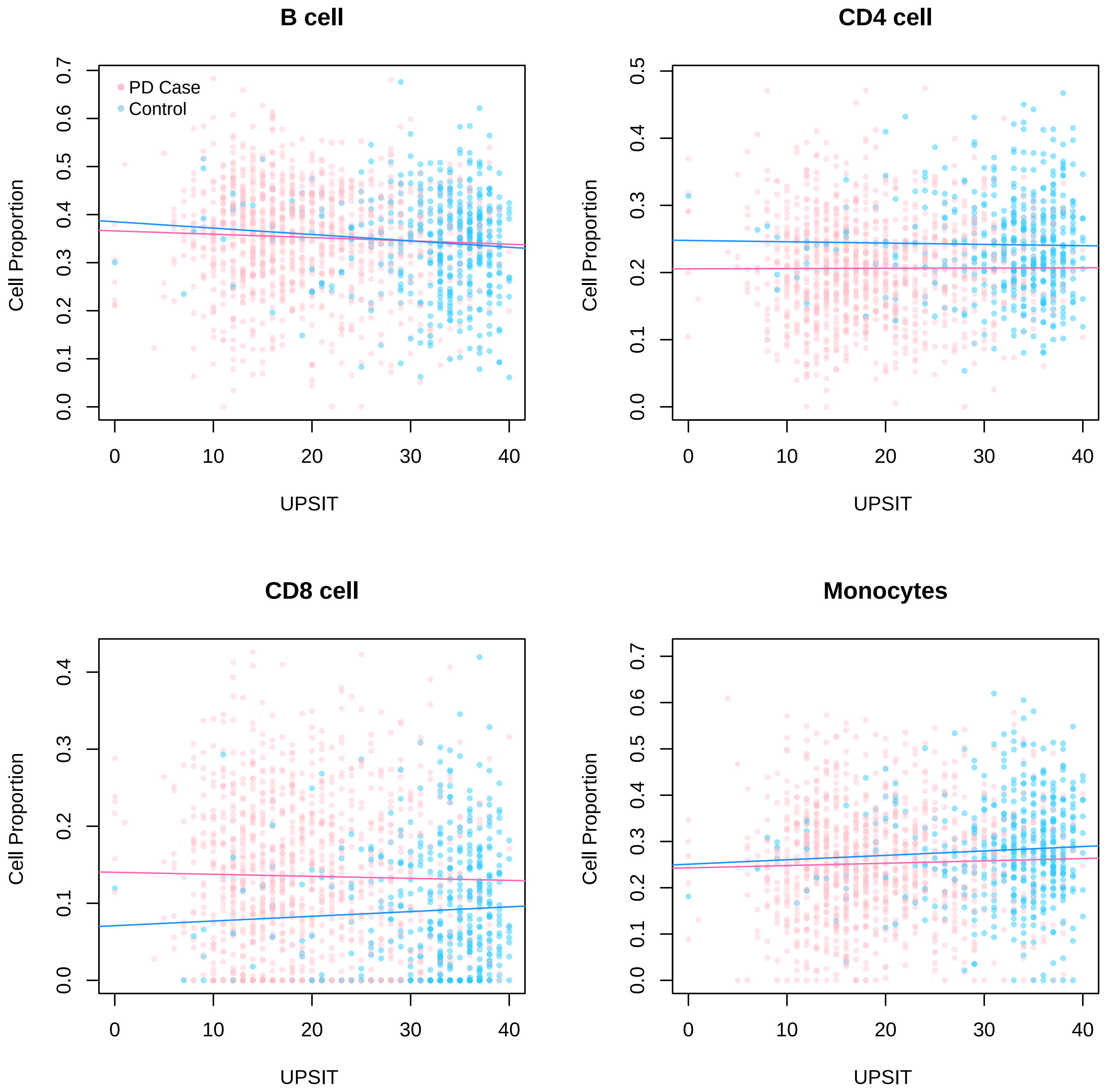

Figure 19: Relationship between total Smell Identification Test Score (UPSIT) and cell type proportion of B cells, CD4 cells, CD8 cells, and Monocytes, respectively. Different colors illustrate PD cases and healthy controls.

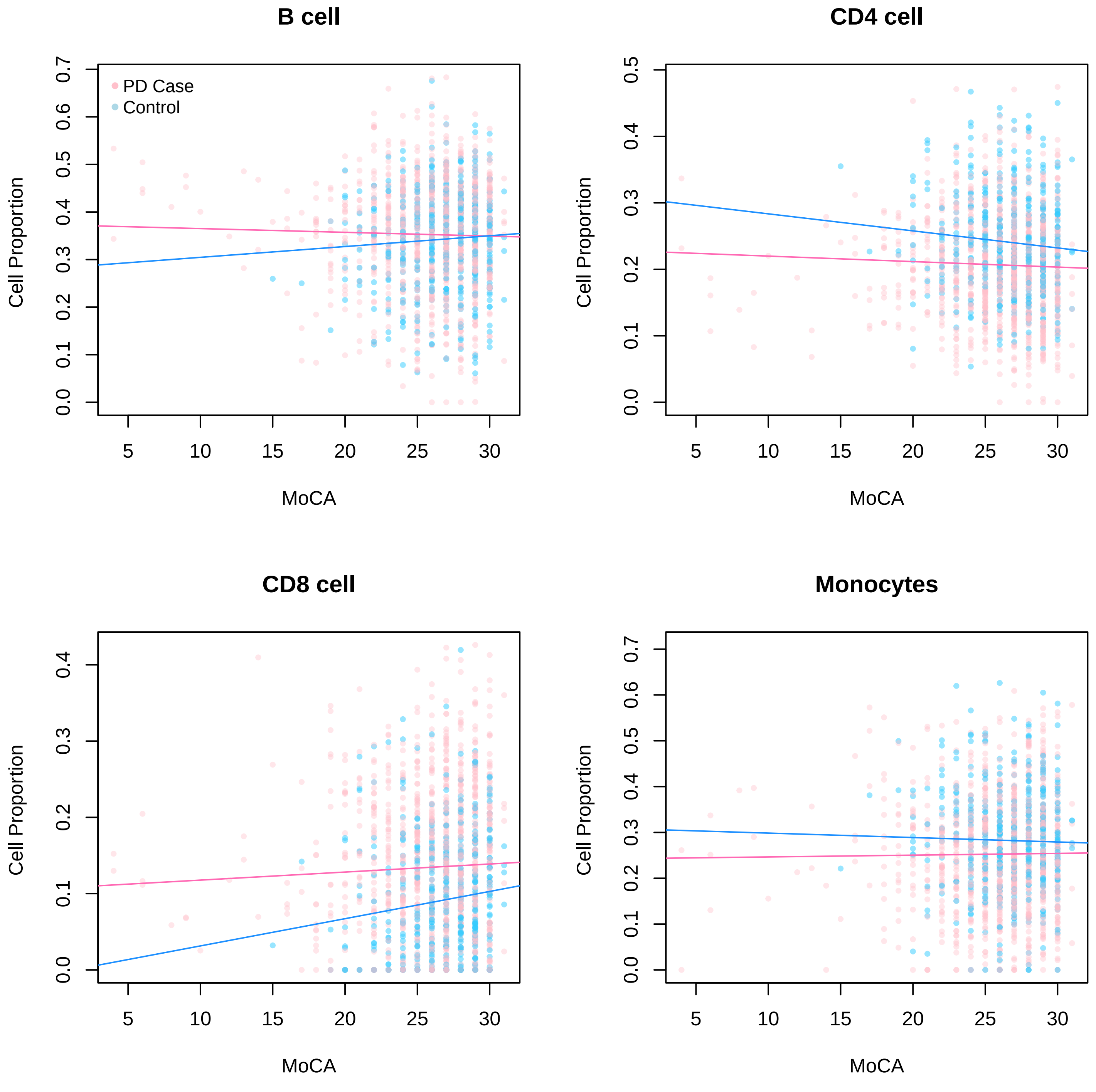

Figure 20: Relationship between total Montreal Cognitive Assessment Score (MoCA) and cell type proportion of B cells, CD4 cells, CD8 cells, and Monocytes, respectively. Different colors illustrate PD cases and healthy controls.

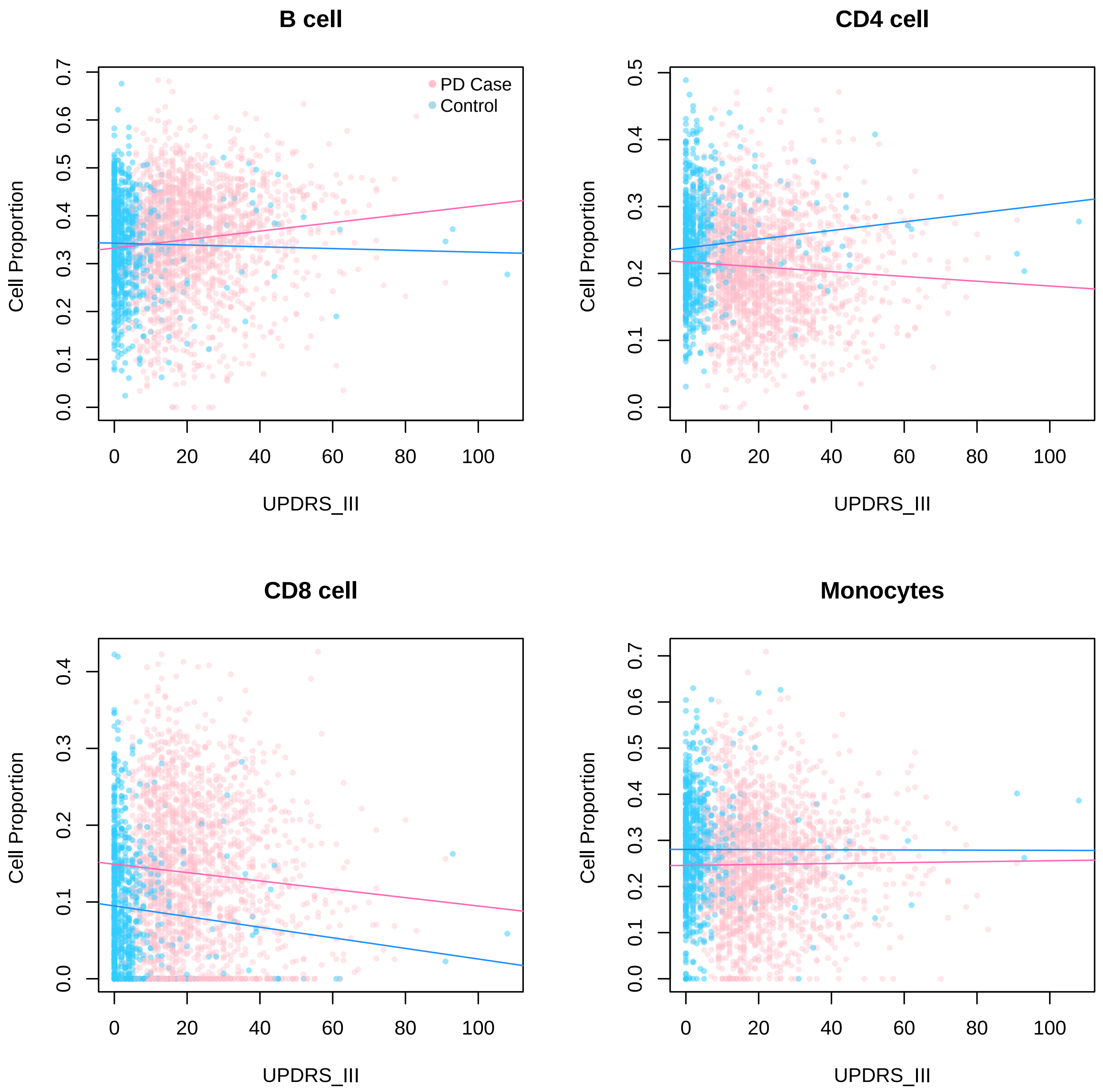

Figure 21: Relationship between total Unified Parkinson's Disease Rating Scale Part III (UPDRS III) and cell type proportion of B cells, CD4 cells, CD8 cells, and Monocytes, respectively. Different colors illustrate PD cases and healthy controls.

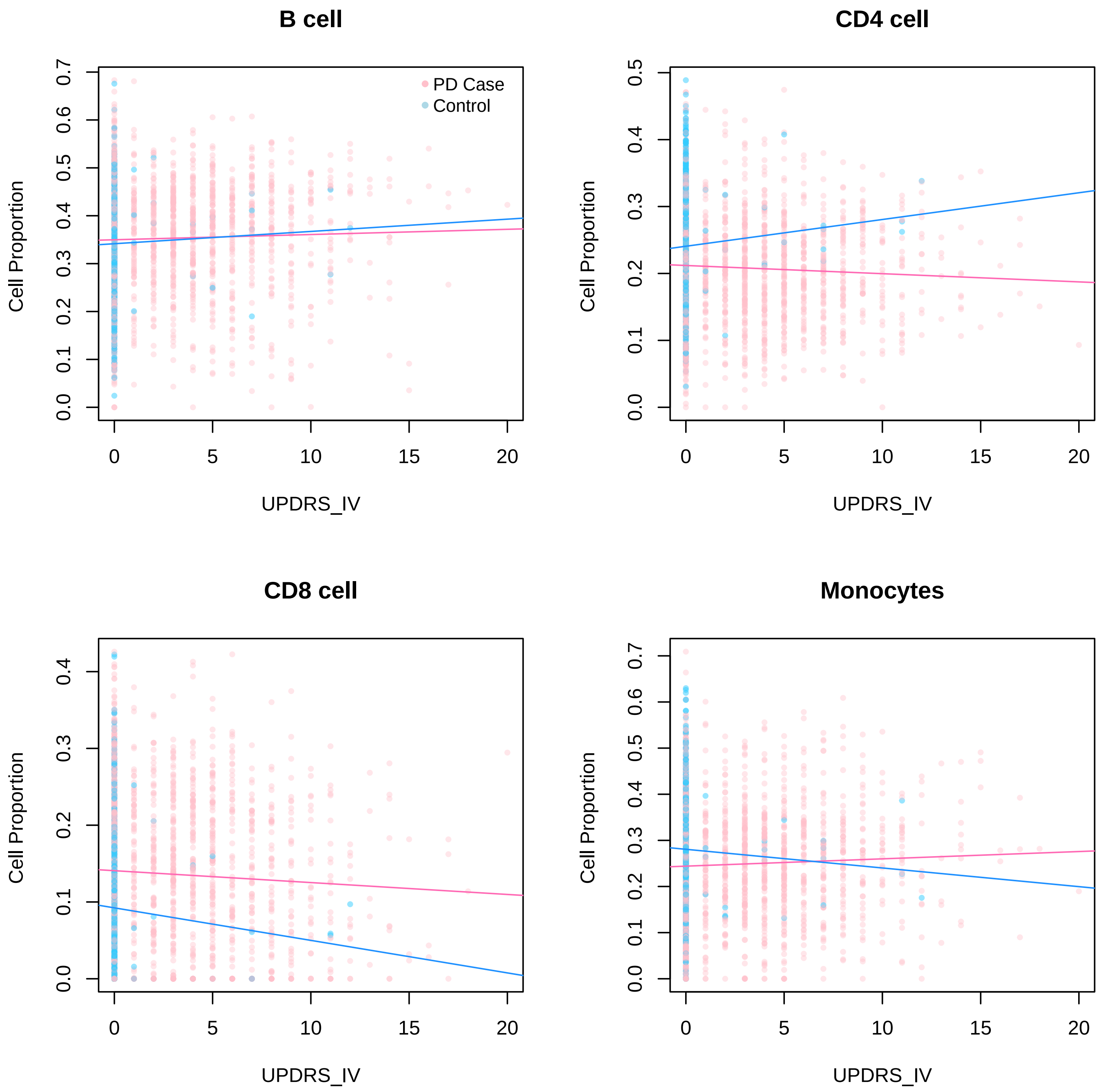

Figure 22: Relationship between total Unified Parkinson's Disease Rating Scale Part IV (UPDRS IV) and cell type proportion of B cells, CD4 cells, CD8 cells, and Monocytes, respectively. Different colors illustrate PD cases and healthy controls.

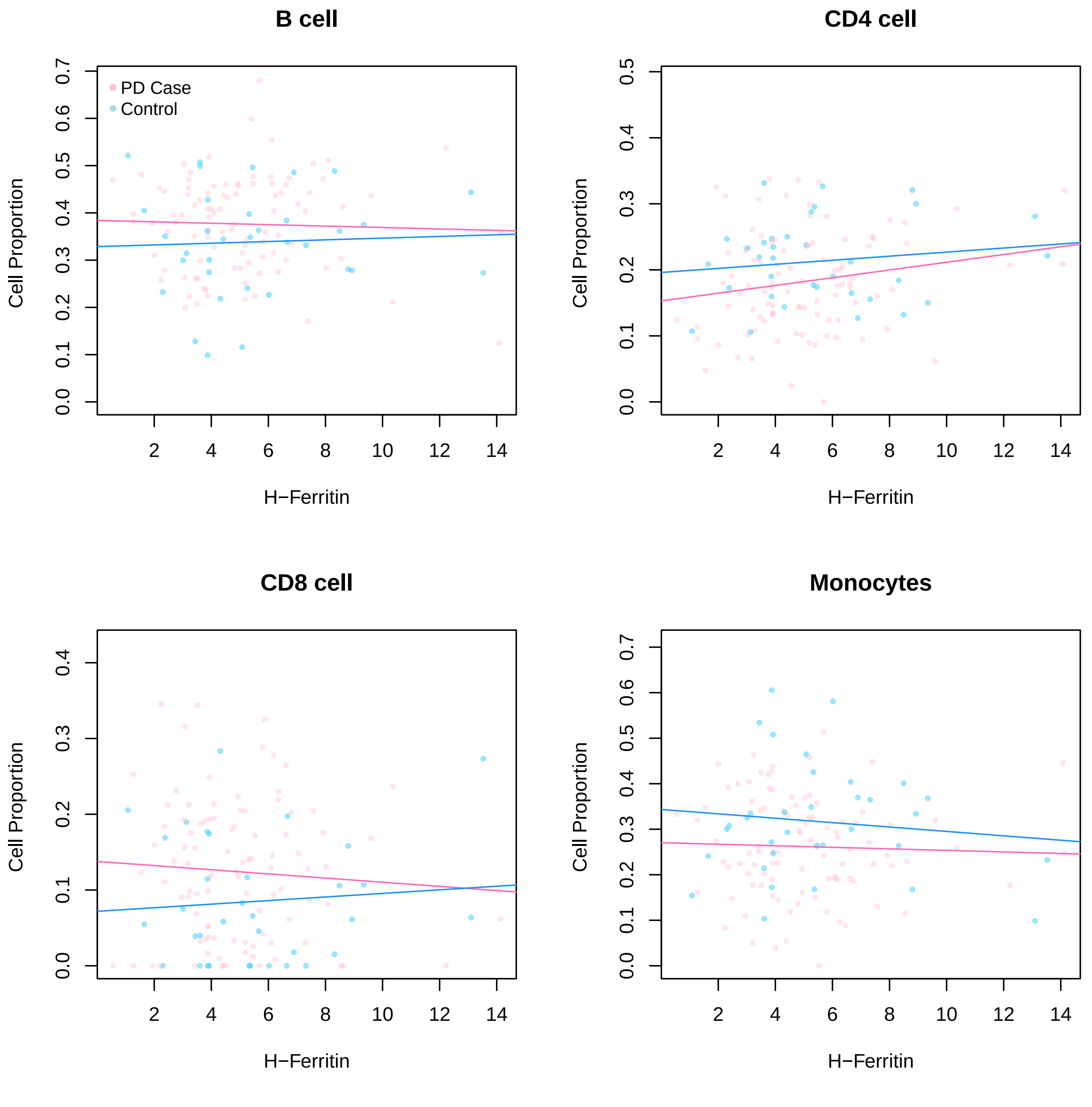

Figure 23: Relationship between Cerebrospinal fluid ferritin levels (H-Ferritin) and cell type proportion of B cells, CD4 cells, CD8 cells, and Monocytes, respectively. Different colors illustrate PD cases and healthy controls.

##### 4.2 TEDDY: The Environmental Determinants of Diabetes in the Young.

This plots shows that the NK cell proportions, solved by CIBERSORT, among participants who developed IA at a young age compared to controls. There is no discernible difference in the proportion trends between cases and controls.

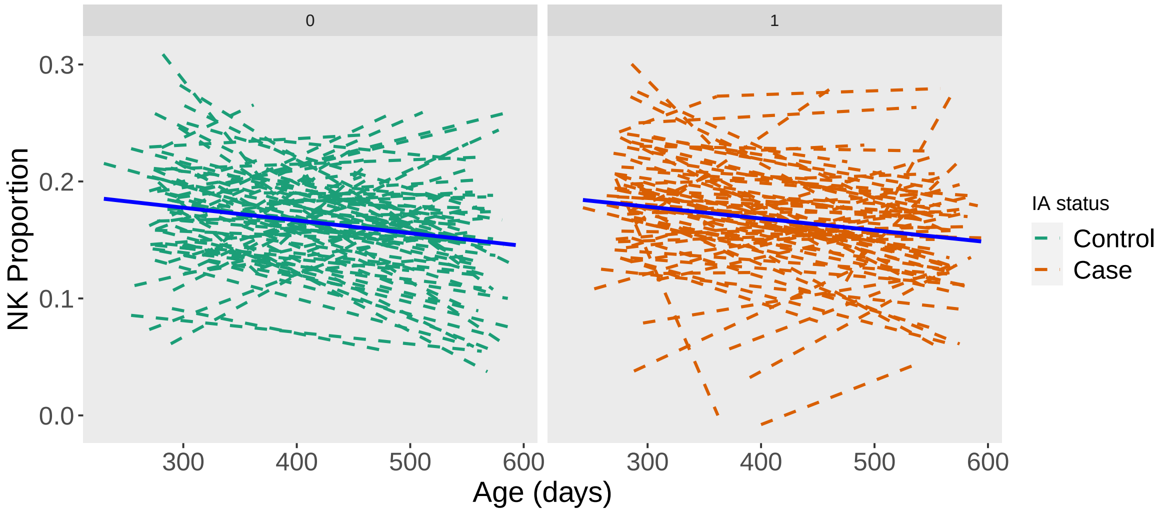

Figure 24: NK cell proportions along infant's age (in days) at sample collection, by case and control status. Average fitted lines (solid) overlay individual-specific lines (dashed).
